# A dominant-negative variant in the dopamine transporter PDZ-binding motif is linked to parkinsonism and neuropsychiatric disease

**DOI:** 10.1101/2020.10.12.336461

**Authors:** Freja Herborg, Kathrine L. Jensen, Sasha Tolstoy, Natascha V. Arends, Leonie P. Posselt, Aparna Shekar, Jenny Aguilar, Viktor K. Lund, Kevin Erreger, Mattias Rickhag, Matthew D. Lykas, Markus N. Lonsdale, Troels Rahbek-Clemmensen, Andreas T. Sørensen, Amy H. Newman, Annemette Løkkegaard, Ole Kjærulff, Thomas Werge, Lisbeth B. Møller, Heinrich JG Matthies, Aurelio Galli, Lena E. Hjermind, Ulrik Gether

## Abstract

Dopaminergic dysfunction is central to movement disorders and mental diseases. The dopamine transporter (DAT) is essential for the regulation of extracellular dopamine but the genetic and mechanistic link between DAT function and dopamine-related pathologies remains elusive. Particularly, the pathophysiological significance of monoallelic missense mutations in DAT is unknown. Here we identify a novel coding DAT variant, DAT-K619N, in a patient with early-onset parkinsonism and comorbid neuropsychiatric disease and in 22 individuals from exome-sequenced samples of neuropsychiatric patients. The variant localizes to the critical C-terminal PDZ-binding motif of DAT and causes reduced uptake capacity, decreased surface expression, and accelerated turnover of DAT *in vitro*. *In vivo*, we demonstrate that expression of DAT-K619N in mice and *dropsophila* imposes impairments in dopamine transmission with accompanying changes in dopamine-directed behaviors. Importantly, both cellular studies and viral overexpression of DAT-K619N in mice show that DAT-K619N has a dominant-negative effect which collectively implies that a single dominant-negative genetic DAT variant can confer risk for neuropsychiatric disease and neurodegenerative early-onset parkinsonism.

## Introduction

Dopamine (DA) exerts strong effects on human behavior by supporting motor initiation and exploration, and through modulation of higher cognitive functions such as reinforcement learning and motivation (Berke, 2018; Iversen & Iversen, 2007). Disturbances in dopaminergic neurotransmission are widely implicated both in neurological diseases, such as Parkinson’s disease (PD), and in psychiatric disorders including attention deficit hyperactivity disorder (ADHD), schizophrenia, bipolar disorder, and depression (Cousins *et al*, 2009; Del Campo *et al*, 2011; Dichter *et al*, 2012; Howes & Kapur, 2009; Iversen & Iversen, 2007; Robison *et al*, 2020). As a critical regulator of dopaminergic neurotransmission (Kristensen *et al*, 2011), the DA transporter (DAT) gene *SLC6A3*, has received enduring attention in candidate gene studies of both PD and psychiatric diseases (Gatt *et al*, 2015). DAT mediates high-affinity reuptake of DA and thereby regulates the spatiotemporal propagation of DA transients and ensures a synthesis-independent DA source to dopaminergic terminals (Sulzer *et al*, 2016). The important role of DAT in exerting behaviorally relevant control of DA homeostasis is substantiated by the acute and long-term effects both of psychostimulant therapeutics and of recreational drugs, such as amphetamine, methylphenidate, and cocaine, which all target DAT.

Although candidate gene studies of PD and mental disorders have reported associations with *SLC6A3* (Gatt *et al.*, 2015; Gizer *et al*, 2009; Zhai *et al*, 2014), genome-wide association studies (GWAS) have not confirmed the existence of disease-associated common *SLC6A3* variants (MacArthur J, 2017). However, several lines of evidence suggest that rare variants of larger effect size may link DAT dysfunction to both parkinsonism and neuropsychiatric disease. Most importantly, a causal relationship between DAT dysfunction and human disease was established with the description of the DAT deficiency syndrome (DTDS) in which inactivating mutations in *SLC6A3* gives rise to a recessively inherited form of infantile parkinsonism-dystonia (Kurian *et al*, 2009). DTDS typically presents in early infancy (Kurian, 1993; Kurian *et al*, 2011; Ng *et al*, 2014); however, we recently identified an adult patient, who carries compound heterozygote missense mutations in *SLC6A3* that only partially disrupt DAT function, and who suffers from an atypical form of DTDS with adult-onset of motor symptoms and comorbid neuropsychiatric disease (Hansen *et al*, 2014). The identification of this patient adds to several reports on heterozygous carriers of coding variants in *SLC6A3* that have been diagnosed with neuropsychiatric disease including bipolar disorder, ADHD, and autism spectrum disorder (ASD) (Bowton *et al*, 2014; Campbell *et al*, 2019; Cartier *et al*, 2015; Grunhage *et al*, 2000; Hamilton *et al*, 2013; Mazei-Robison *et al*, 2005; Sakrikar *et al*, 2012). Most of these DAT variants show deficits in DAT properties and/or trafficking, and several cause behavioral changes *in vivo*, at least when homozygously expressed (Campbell *et al.*, 2019; Cartier *et al.*, 2015; DiCarlo *et al*, 2019; Hamilton *et al.*, 2013; Hansen *et al.*, 2014; Herborg *et al*, 2018; Mazei-Robison *et al.*, 2005; Mergy *et al*, 2014; Sakrikar *et al.*, 2012; Wu *et al*, 2015). Moreover, *SLC6A3* shows a high degree of conservation and is constrained against both ‘loss of function’ variants and missense variants – supporting that perturbations of DAT function may contribute to malfunctional states that impose the negative selection against such variants (Lek *et al*, 2016).

Here, we investigate the mechanistic and functional impact of a recurring rare coding DAT variant, DAT-K619N on dopaminergic neurotransmission. We first present two unrelated patients that carry the DAT-K619N variant. The first patient was identified in the Simon’s Simplex Collection of ASD families (Fischbach & Lord, 2010); the second patient is an adult male with early-onset parkinsonism and comorbid neuropsychiatric disease, for whom we use Single Photon Emission Computerized Tomography (SPECT) scans to reveal a progressive loss of DAT availability. By capitalizing on a large population-based exome sequencing dataset we next uncover that the DAT-K619N mutation is present in additional 21 patients with neuropsychiatric disease, as well as in five control subjects, with a nominally significant association to bipolar disorder. Importantly, the DAT-K619N mutation is located in a C-terminal <u>P</u>SD-95/<u>D</u>iscs-large/<u>Z</u>O-1 homology (PDZ) binding sequence, which is critical for DAT surface expression *in vitro* (Torres *et al*, 2001) and for striatal expression and surface stability *in vivo* (Rickhag *et al*, 2013). We show that the DAT-K619N variant is subject to accelerated cellular turnover that in turn causes impaired surface expression and reduced DA uptake capacity. This functional impairment is dominant-negative towards the WT transporter as supported both by *in vitro* experiments and by viral overexpression in mice. In mice, we also find that unilateral overexpression of DAT-WT and DAT-K619N in nigrostriatal neurons causes diverging effects on DA-directed behavior and, by expressing the DAT-K619N variant in *Drosophila*, we demonstrate that the mutant can drive dopaminergic dysfunction with associated alterations in locomotion. Collectively, our study substantiates the putative role of rare coding DAT variants as genetic risk factors in neuropsychiatric disease, and lend further support to a link between genetic insult to DAT function and neurodegenerative parkinsonism in adults.

## Results

### Identification of the DAT-K619N variant in two unrelated patients with early-onset parkinsonism and/or neuropsychiatric disease

In two unrelated patients from independent samples, we identified the same single nucleotide substitution (c.1857G>C) in exon 15 of *SLC6A3*, giving rise to the coding variant DAT-K619N. The first patient belonged to the Simon Simplex Collection comprising more than 2000 ASD families (Fischbach & Lord, 2010), the second was identified in a previously described cohort of 91 patients with symptoms of early-onset parkinsonism, dystonia, or other unclassified movement disorders (Hansen *et al.*, 2014). Interestingly, the K619 residue is part of a PDZ-binding domain, that has been reported to regulate DAT trafficking and to be essential for normal expression of DAT at striatal release sites (Bjerggaard *et al*, 2004; Rickhag *et al.*, 2013; Torres *et al.*, 2001) (Figure 1A). Only patient 2 was available for further clinical evaluation. He is a Caucasian male (aged 41 years at the time of referral) diagnosed with early-onset parkinsonism and neuropsychiatric disease. Inheritance investigations showed that the DAT-K619N allele was paternally transmitted (Figure 1B), but it is not known if the father has neurological or neuropsychiatric symptoms or signs, as he did not want to be examined. According to clinical records, patient 2 reported insidious onset of right-sided hand tremor in his late thirties. His symptoms were initially interpreted as being part of a psychiatric disorder and autism was suspected, but he was diagnosed with early-onset parkinsonism when his motor symptoms worsened. At age 41, he presented with unequivocal hemiparkinsonism with right-sided symptoms. A brain MRI scan at age 39 was reportedly normal, except for the presence of more, small (2-4 mm) white matter lesions than expected for this age. Electroencephalography, as well as visual and motor evoked potentials, were normal. Genetic testing for selected relevant spinocerebellar ataxias, Huntington’s disease-like syndromes, and hereditary parkinsonism uncovered no extended trinucleotide repeat in *ATXN1-3, CACNA1A*, *TBP*, and *FMR1*, no pathogenic variants in *PRKN* and *GCH1*, and no hotspot variant in LRRK2 (Gly2019Ser) or common point variants and copy number variations in *DJ1, PINK, UCHL1, SNCA*, and *ATP13A2*. Further, no pathogenic variants were found in *NPC1, NPC2*, or *NOTCH3.* Patient 2 was not formally assessed by specialists in child psychiatry during his childhood and adolescence. As an adult, however, he has had several episodes of self-injury and has been in intermittent contact with in-patient and out-patient psychiatric facilities. He was diagnosed with an unspecified personality disorder with evasive and schizophrenic traits (ICD-10-CM code F60.9) and periodic depression (F33.9) at age 43. Treatment of parkinsonian symptoms were initially attempted with DA agonists in low-to moderate doses (with some positive effects, but also side-effects), and levodopa/carbidopa was later administered with a positive effect. Moreover, the patient has received treatments to improve his psychiatric symptoms including risperidone in low dose (discontinued due to side effects), and escitalopram in a shorter period. All treatments were with variable compliance.

**Figure 1.**
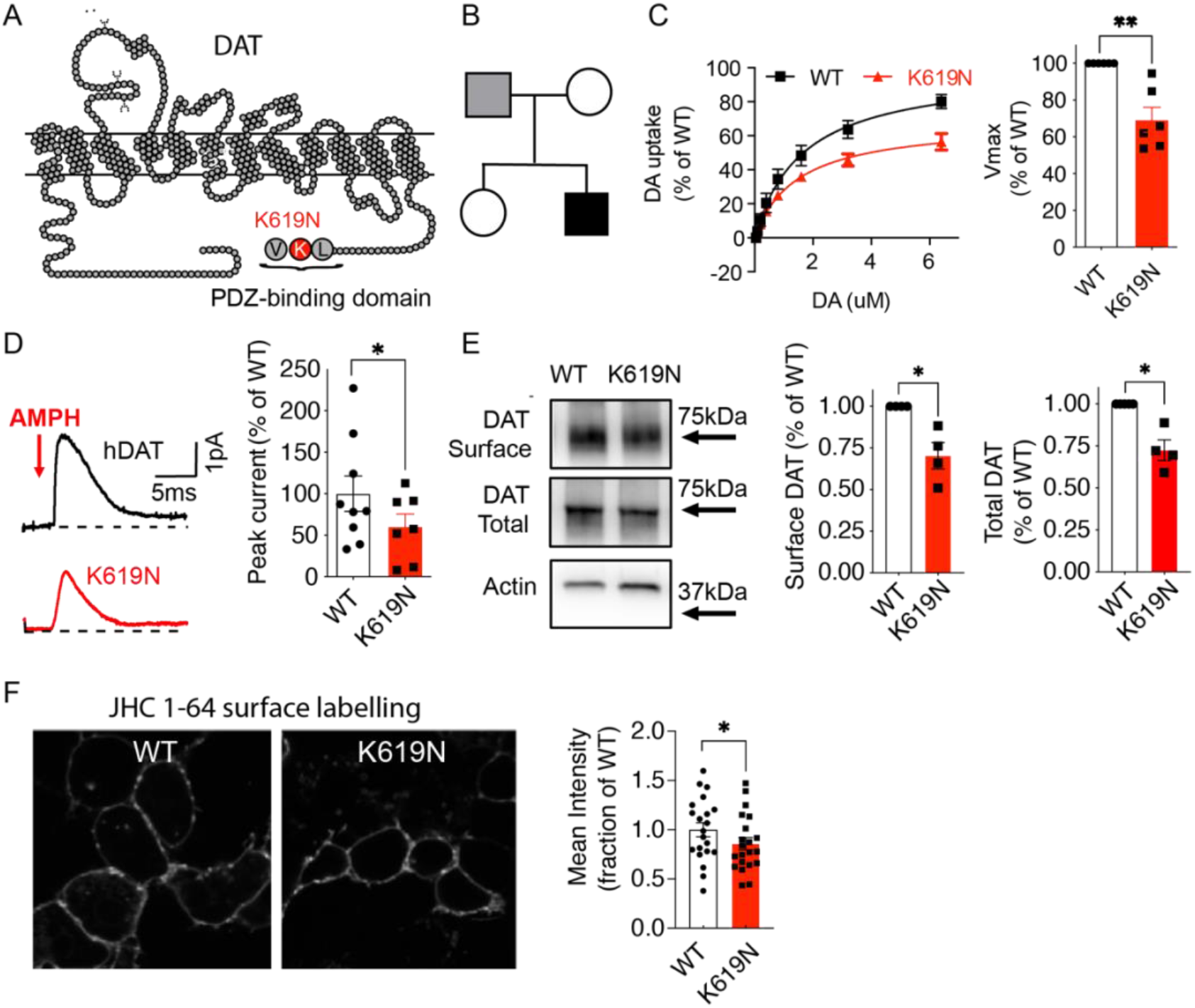
DAT-K619N displays functional impairments and reduced surface expression *in vitro*. (A) Snake diagram of DAT demonstrating the C-terminal location of the DAT-K619N mutation within a PDZ-binding domain. (B) Family tree with the index patient shown in black shading. Only the patient’s parents were available for genetic analysis, which revealed paternal transmission of the DAT-K619N allele. It is unknown if the father has neurological or psychological symptoms. (C-F) Evaluation of DAT-K619N functions and surface expression in transiently transfected HEK293 cells. (C) Functional comparison of DAT-K619N to WT using [^3^H]-dopamine (DA) uptake. Uptake curves (left) are average curves of six experiments each performed in triplicates and normalized to the fitted maximal uptake capacity (V_max_) of DAT-WT. The DAT-K619N variant demonstrates reduced V_max_ (right bar diagram) compared to DAT-WT (p<0.01, one-sample t-test, N=6) with no accompanying change in K_M_ (K_M_=1.4±0.3 μM for K619N vs. 1.7±0.4 μM for DAT-WT, p>0.05 Student’s t-test). (D) Amphetamine (AMPH)-induced amperometric currents with representative traces of amperometric currents (left) and quantification of the amphetamine-induced peak currents relative to DAT-WT (right). DA-release through DAT-K619N is impaired compared to DAT-WT (p<0.05, one-sample t-test, N=9 for WT and N=7 for DAT-K619N). (E) Surface biotinylation of transiently transfected HEK293 cells. The amount of DAT-K619N relative to DAT-WT is decreased in both the surface and the total protein fractions (p<0.05, one-sample t-test, N=4). (F) Confocal live imaging of surface expressed WT and DAT-K619N in HEK239 cells by labelling with the fluorescent cocaine analogue, JHC 1-64 (20 nM). Mean intensity of the JCH 1-64 signal is reduced for DAT-K619N relative to DAT-WT. Images were acquired from four independent transfections and intensities were normalized to WT mean intensity for each imaging session (p<0.05, one-sample t-test, N=21 images per group). Data are means±SEM. *p<0.05, **p<0.01.

Patient 2 is currently 53 years and presents with general bradykinesia, severe rigidity, resting and intention tremor of the upper extremities without consistent sidedness of symptoms (Supplementary video 1). The tremor is slightly more irregular than the usual parkinsonian tremor, probably due to dystonic elements. His gait is parkinsonian with small shuffling steps, kyphosis and severe difficulties in turning around (Supplementary video 2). Postural instability is obvious, and retropulsion pull test is not possible as he tends to fall spontaneously. He has severe oligomimia, but his eye movements show normal pursuit with complete range and no square wave jerks. Saccades are normal, including initiation, velocity and range. His voice is slightly hoarse, but not severely hypophonic (Supplemental video 2). Current treatment is Sinemet (Levodopa and carbidopa) (900mg/day). He has been offered, but declined advanced treatment (continued duodenal infusion of levodopa/carbidopa (DuoDopa ®)).

### Population genetics of the DAT-K619N variant

Like other previously identified DAT coding variants (Herborg *et al.*, 2018; Lek *et al.*, 2016), DAT-K619N (rs200712598) can be found in genetic databases of the general population. With a reported frequency of only 0.00023 (gnomAD database (Karczewski *et al*, 2019)), DAT-K619N is still among the most common missense variants in *SLC6A3* (Karczewski *et al.*, 2019; Lek *et al.*, 2016). Since the two identified patients carrying the DAT-K619N variant both suffer from neuropsychiatric disease, we also evaluated the occurrence of DAT-K619N in 19,851 exome-sequenced samples from the integrated psychiatric research (iPSYCH) consortium case-cohort covering five major mental disorders: ADHD, ASD, schizophrenia, depression, and bipolar disorder (Pedersen *et al*, 2018). All subjects of the study sample were born between May 1, 1981, and December 31, 2005, with a follow-up period from 1994 to 2016. Following sample and variant quality control (see supplementary methods for details), 17,339 samples were included in the analysis, comprising 4885 controls and 12327 cases diagnosed with at least one of the following diseases: ADHD (N=4548), ASD (N=4937), schizophrenia (N=3052), single or recurrent depression (N=2623), or bipolar disorder (N=1313) (Table 1). The K619N variant was observed in 26 subjects: 5 controls and 21 cases. A carrier-based association analysis did not establish a disease association across all diagnostic groups (P=0.29, odds ratio (OR) [95%CI]: 1.7 [0.68-4.3]). Analysis of individual diagnostic groups, however, did show a general trend of increased odd ratios across all diagnostic categories and demonstrated a nominally significant association with bipolar disorder (Table 1). Of notice, the iPSYCH sample includes relatively young individuals and later diagnosis, re-diagnosis, or additional diagnosis may occur in the future, but the median and mean age of DAT-K619N carriers among cases and controls were comparable (case-carriers: mean age=25.68 years, median age=25.42 years, StdError=0.98; control-carriers: mean age=28,75 years, median age=27,36 years, StdError=1.73).

**Table 1:** DAT-K619N carriers in the iPSYCH study sample. Exome sequencing from the first phase of the iPSYCH case-cohort (described in (Pedersen *et al.*, 2018)), covering five major neuropsychiatric diagnosis: ADHD, ASD, schizophrenia, depression, and bipolar disorder covered by the denoted ICD10 diagnoses. Identification was achieved through linkage to the Danish Central Research Register, which at the time of linkage contained information about all contacts up until 2016. Abbreviations: ADHD, attention-deficit/hyperactivity disorder; ASD, autism spectrum disorder, ASD; The ICD10, International Classification of Diseases, 10th revision; iPSYCH, Integrative Psychiatric Research; QC, quality control.

| Sample<sup>a</sup> | Initial sample size | Sample size post QC | ICD10 diagnosis | Follow up period | # LoF carriers | P-value | OR (95% CI) |
| --- | --- | --- | --- | --- | --- | --- | --- |
| Controls | 5554 | 4885 | Randomly sampled | 1994-2016 | 5 |  |  |
| Any case | 14283 | 12454 | ICD codes listed below | 1994-2016 | 21 | 0.39 | 1.7<br>(0.64-4.1) |
| ADHD | 5040 | 4549 | F90.0 | 1994-2016 | 8 | 0.41 | 1.7<br>(0.56-4.7) |
| ASD | 5694 | 4938 | F84.0, F84.0, F84.5, F84.8, F84. or F84.9 | 1994-2016 | 8 | 0.58 | 1.6<br>(0.51-4.3) |
| Schizophrenia | 3686 | 3052 | F20 | 1994-2016 | 5 | 0.74 | 1.3<br>(0.39-4.2) |
| Single and recurrent depression | 2967 | 2623 | F32-33 | 1994-2016 | 7 | 0.13 | 2.6.<br>(0.92-7.2) |
| Bipolar disorder | 1528 | 1314 | F30-F31 | 1994-2016 | 5 | **0.041** | **3.7**<br>**(1.2-11.3)** |

<sup>a</sup>Samples are not mutually exclusive

### DAT-K619N variant shows impaired DA uptake, amphetamine-induced efflux, and surface expression

To investigate the molecular phenotype of DAT-K619N, we first assessed DA uptake in transiently transfected HEK239 cells. Compared to DAT-WT, DAT-K619N demonstrated impaired maximal DA uptake capacity (V_max_=69± 7% of DAT-WT) without any change in K_M_ (K_M_=1.4±0.3 μM for DAT-K619N vs. 1.7±0.4 μM for DAT-WT, Figure 1C). DAT is known to also mediate reverse transport of DA in response to amphetamine, which acts as a competitive substrate and promotes an efflux-prone conformation of DAT (Sulzer, 2011). Amperometric recordings on transiently transfected HEK293 cells, loaded with DA and stimulated with 10μM amphetamine, showed reduced amphetamine-induced DA efflux with a peak current for DAT-K619N of ~60% of the WT current (Figure 1D).

Next, we addressed if a reduction in surface expression might be responsible for the functional deficits of DAT-K619N. Indeed, whole-cell surface biotinylation experiments demonstrated a decrease in surface-expressed DAT-K619N compared to DAT-WT (70±8 % of DAT-WT, Figure 1E). The total expression of maturely glycosylated K619N was also reduced (79±1 % of WT, p<0.01), suggesting that the reduced surface level is the result of a reduction in the total cellular level of transporter. The biotinylation data was further supported by assessing surface expression for DAT-K619N using a fluorescent cocaine analogue, JHC 1-64 (20nM for 20 min, RT), which allows selective live labelling of the surface-expressed transporters (Cha *et al*, 2005; Eriksen *et al*, 2009). JHC 1-64 produced a clear membrane labelling of both DAT-WT and DAT-K619N expressing cells, but the mean fluorescent intensity was reduced in cells expressing DAT-K619N compared to DAT-WT (mean JHC 1-64 intensity 86±6% of DAT-WT) consistent with lower surface levels (Figure 1F).

The functional effects of mutating DAT’s C-terminal PDZ-binding sequence can vary considerably between different expression systems (Rickhag *et al.*, 2013). Moreover, we have previously found that disease-associated DAT variants display changes in affinities for ligands and for co-transported Na^+^ and/or Cl^−^ ions, and show alterations in Zn^2+^-dependent regulation of DA uptake in COS-7 cells (Herborg *et al.*, 2018). We therefore evaluated DA uptake, [^3^H]-CFT ([^3^H]-2β-carbomethoxy-3β-(4-fluorophenyl)tropane) binding, ligand affinities, as well as ion coordination and Zn^2+^ regulation in transiently transfected Cos7 cells to further elucidate the molecular phenotype of DAT-K619N (28). As observed in HEK293 cells, the uptake capacity of DAT-K619N was reduced by ~30%, and we observed a comparable reduction in [^3^H]-CFT binding capacity (B_max_=75± 5% of DAT-WT, Supplementary Figure 1). Neither the apparent affinity for DA nor CFT was different between DAT-WT and the DAT-K619N variant and potencies of amphetamine, methylphenidate, and cocaine to inhibit uptake were unaffected for DAT-K619N. Furthermore, DAT-K619N showed no changes in coordination of co-transported Na^+^ and Cl^−^ ions, as assessed by DA uptake measurements at increasing Na^+^ and Cl^−^ concentrations respectively. Likewise, Zn^2+^-dependent regulation of DA uptake was similar for DAT-WT and DAT-K619N (Supplementary Figure 1 and Supplementary Table 1). Summarized, the functional phenotype of DAT-K619N could be recapitulated in COS-7 cells, supporting reduced expression and no conformational perturbations of DAT-K619N.

### Live fluorescent imaging uncovers accelerated turnover-rate and increased lysosomal targeting of DAT-K619N

To obtain further insight into the cellular processing of DAT-K619N, we took advantage of genetically encoded fluorescent timers (FT) that change color from blue to red in a time-dependent manner and thereby permit concurrent spatial and temporal visualization of protein trafficking (Knop & Edgar, 2014; Subach *et al*, 2009). We generated fusion proteins of a blue/red fluorescent timer, with reported slow kinetic properties (SlowFT), attached to the N-terminus of DAT-WT and DAT-K619N, and expressed them in a neuronally-derived catecholaminergic cell line (Cath.a-differentiated, CAD cells) (Qi *et al*, 1997) with relatively large somas, making them suited for live microscopy. Imaging was performed following surface labelling of DAT with an Oregon Green-conjugated fluorescent cocaine analogue, MFZ-9-18 (Eriksen *et al.*, 2009), which enabled identification of transfected cells without directly imaging of the SlowFT (see Methods) and allowed subsequent analysis of surface signal versus intracellularly located DAT. For both SlowFT-DAT-WT and SlowFT-DAT-K619N, the fluorescence from the blue and red timer forms colocalized at the membrane, but showed only a partial overlap for the intracellular punctate structures (Figure 2A). Interestingly, an inspection of the cells indicated a relatively low intensity of the red form of SlowFT-DAT-K619N as compared to SlowFT-DAT-WT. Consistently, quantification of the overall red-to-blue ratio, a relative measure of protein “age”, was reduced for SlowFT-DAT-K619N compared to SlowFT-DAT-WT, indicating that the expressed DAT-K619N on average is “younger” than DAT-WT (Figure 2B, C). Next, we used the signal of the MFZ-9-18 cocaine analogue bound to DAT in the plasma membrane to create a mask that enabled us to isolate the surface compartment thus to analyze the red-to-blue ratios or ‘relative ages’ and for surface- and intracellularly localized transporters separately. This revealed a reduction in the mean red-to-blue ratio of SlowFT-DAT-K619N compared to SlowFT-DAT-WT protein in both compartments (Figure 2C). Of notice, we also found that the mean red-to-blue ratio of SlowFT-DAT-WT in the ‘surface/MFZ-9-18 positive compartment’ was higher than the red-to-blue ratio of the intracellular compartment, indicating that surface expressed SlowFT-DAT-WT, on average, is older than that found intracellularly. By contrast, the mean red-to-blue ratio for SlowFT-DAT-K619N in the surface compartment was smaller than mean ratio for the intracellular compartment (Supplementary Figure 2), supporting that the cellular processing of SlowFT-DAT-K619N differs from SlowFT-DAT-WT. The overall reduction in mean age of DAT-K619N compared to DAT-WT, as well as the increase in mean age of DAT-K619N residing in intracellular compartments relative to that on the surface is consistent with an accelerated turnover rate of DAT-K619N. We further rationalized that an accelerated turnover of DAT-K619N likely would be associated with enhanced lysosomal degradation. Accordingly, we expressed mCherry-tagged versions of DAT-WT and DAT-K619N in CAD cells and concurrently visualized the lysosomes with lysotracker. Quantification of the mCherry mean intensity confirmed a reduction for mCherry-DAT-K619N compared to mCherry-DAT-WT (Supplementary Figure 3). Using Manders coefficients to quantify the fraction of the total mCherry signal that overlapped with lysosomes we found that the fractional overlap between mCherry-DAT-K619N and lysotracker-positive compartments was significantly larger (~33%) than for mCherry-DAT-WT (fractional overlap=0.060±0.006 for DAT-WT vs 0.080±0.007 for DAT-K619N, Supplementary Figure 3), supporting that the DAT-K619N variant may be subject to enhanced lysosomal degradation.

**Figure 2.**
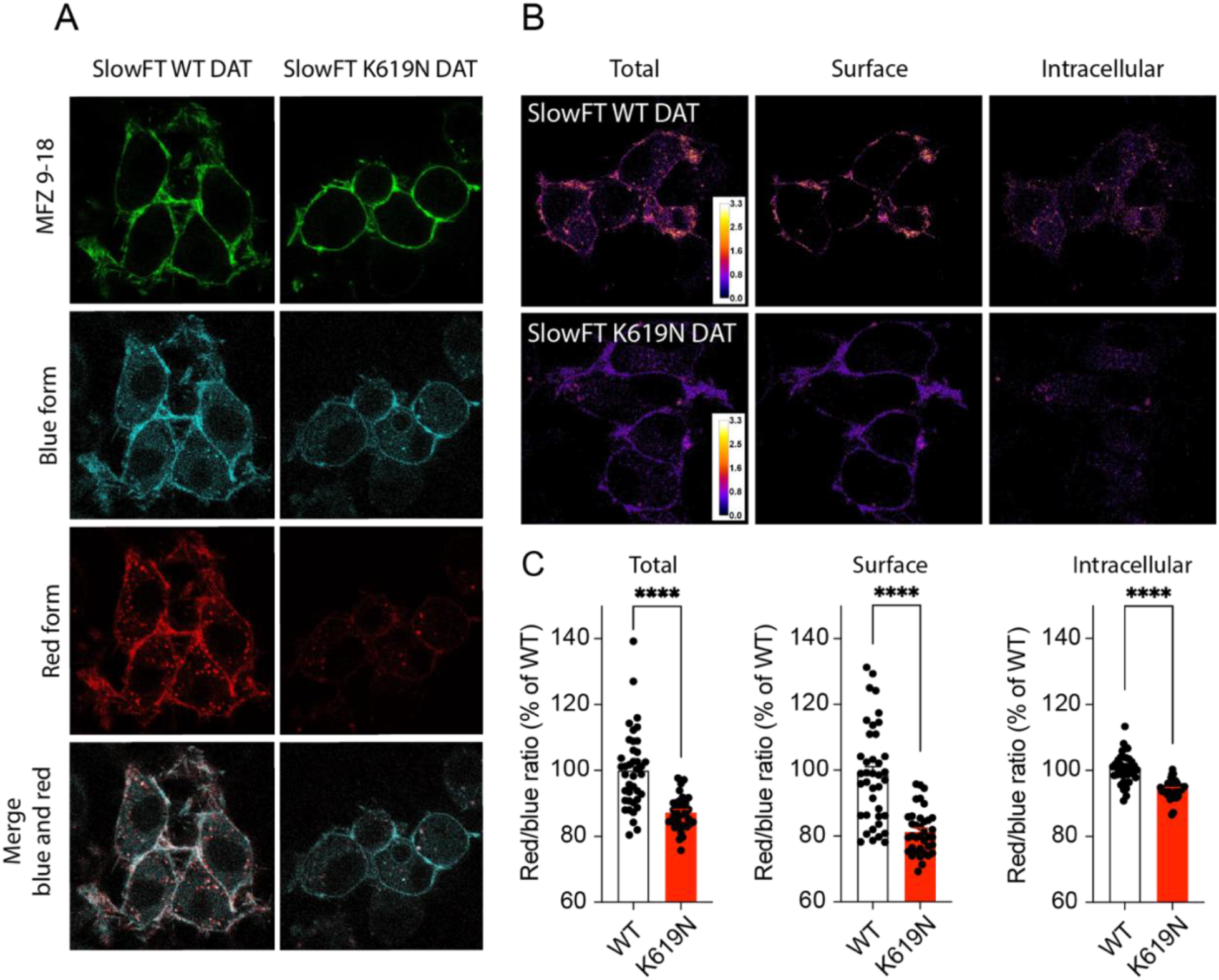
DAT-K619N shows altered cellular processing. Spatiotemporal visualization of DAT-WT and DAT-K619N in transfected CAD cells, using N-terminal tagging with the fluorescent timer, SlowFT, that changes color from blue to red over time (Subach *et al.*, 2009). (A) Images from live confocal microscopy of SlowFT-DAT-WT and SlowFT-DAT-K619N. A green fluorescent cocaine analogue, MFZ-9-18 was used to identify transfected cells and as a surface indicator in postimaging analysis of the surface and intracellular fractions. The blue and red timer forms are shown individually and merged. Note the apparent reduction in the red form of SlowFT-DAT-K619N (B) Pseudo-color images of the red-to-blue ratios, which is a relative measure of age. Warm colors indicate older protein (higher red-to-blue ratios). (C) Quantification of mean red-to-blue ratios for SlowFT-DAT-WT and SlowFT-DAT-K619N, normalized to the mean red-to-blue ratio of SlowFT-DAT-WT for each imaging session. SlowFT-DAT-K619N is on average younger than SlowFT-DAT-WT, both for the total protein and in the surface and intracellular compartments, consistent with an accelerated turnover of SlowFT-DAT-K619N compared to SlowFT-DAT-WT (p<0.0001, one-sample t-test, N=36-39). Data are shown as means ± SEM. Images are representative of 36-39 images from three independent experiments. Image analysis was done in ImageJ (see methods) ****p<0.0001.

**Figure 3.**
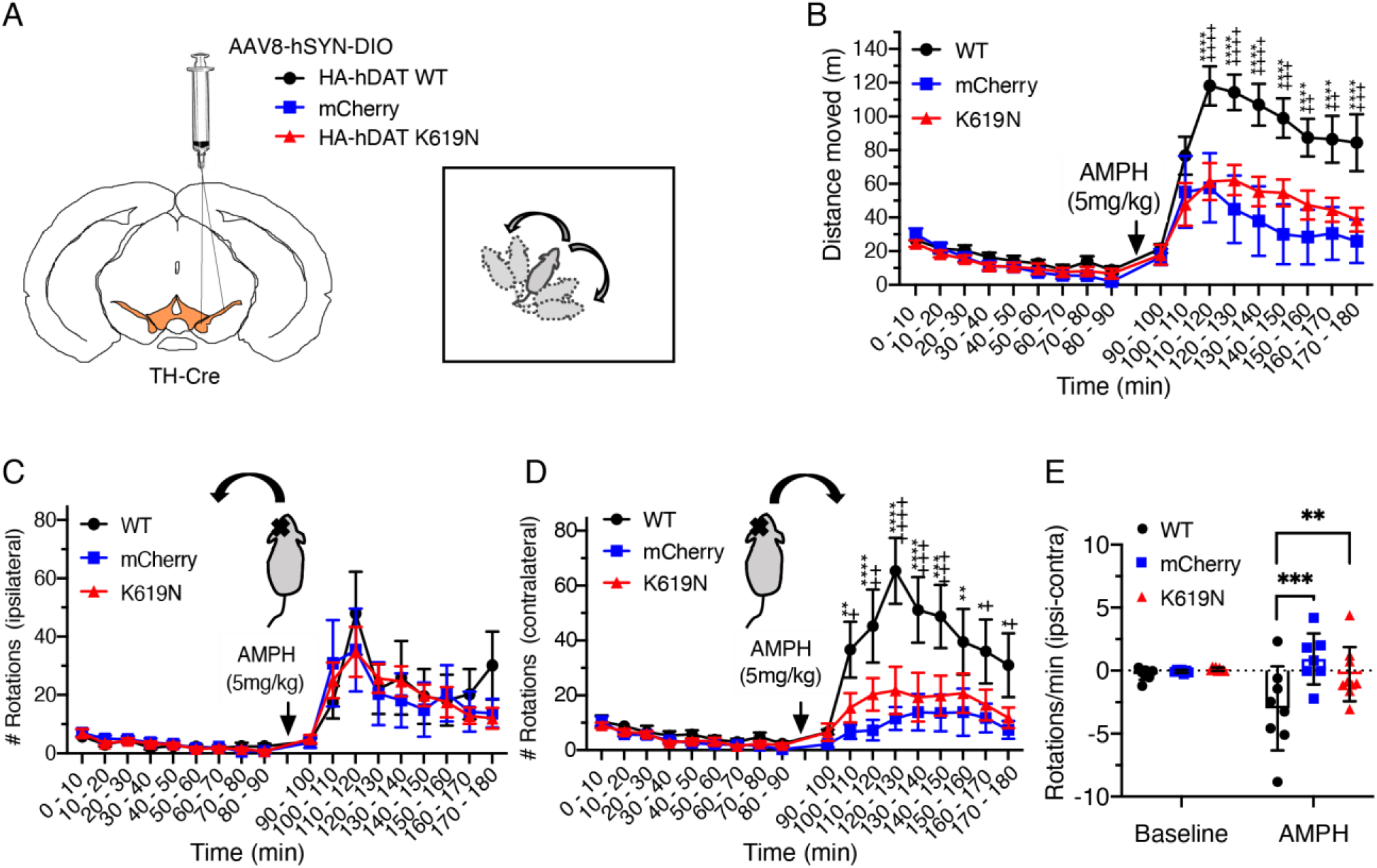
AMPH-induced rotations reveal differential DA-controlled behaviors following unilateral expression of DAT-WT and DAT-K619N. The effect of HA-DAT-WT and HA-DAT-K619N on striatal DA homeostasis was evaluated by unilateral expression and open-field assessment of AMPH-induced rotations. (A) TH-Cre mice were injected unilaterally in substantia nigra with AAV encoding HA-DAT-WT, HA-DAT-K619N or mCherry, and amphetamine (AMPH)-induced rotations were evaluated in the open-field setup three weeks after injections. (B) Distance travelled in open-field chambers 90 min before and after AMPH injections (5 mg/kg i.p.). Mice injected with HA-DAT-WT differ markedly from HA-DAT-K619N injected mice by showing enhanced activity (repeated measures 2-way ANOVA followed by Holm-Sidak multiple comparisons test, N=7-9 mice). (C) Evaluation of ipsilateral rotations show no difference between HA-DAT-WT, HA-DAT-K619N, or mCherry injected mice upon AMPH exposure (p>0.05, repeated measures 2-way ANOVA). (D) The number of contralateral rotations is enhanced in mice expressing HA-DAT-WT following AMPH treatment, supporting that the effect of HA-DAT-K619N differs markedly from HA-DAT-WT *in vivo.* (E) rotational laterality assessed as ‘ipsilateral-contralateral rotations’ per min before and after amphetamine injections. Only HA-DAT-WT injected mice display lateralized rotational behavior (2-way ANOVA followed by Holm-Sidak multiple comparisons test, N=7-9 mice). In panel B and D, crosses are used to mark statistical differences between HA-DAT-WT and HA-DAT-K619N and stars are used to mark statistical differences between HA-DAT-WT and mCherry. No differences were found between HA-DAT-K619N and mCherry. ****/++++ p<0.0001, ***/+++ p<0.001, **/++p<0.01, */+p<0.05). All data are shown as means ± SEM.

### Unilateral nigral overexpression of K619N does not generate rotational bias in response to AMPH

To directly compare effect of DAT-K619N on striatal dopamine function with that of DAT-WT *in vivo*, we expressed DAT-K619N or DAT-WT unilaterally in substantia nigra and determined how these transgenes affected in an open-field test AMPH-induced (5mg/kg) rotations, which is a sensitive measure for asymmetry in striatal DA release (Bay Konig *et al*, 2019; Bjorklund & Dunnett, 2019).. To enable specific detection of the virally encoded transgenes, an HA-tag was fused to the N-terminal of DAT-WT (HA-DAT-WT) and DAT-K619N (HA-DAT-K619N). The resulting adeno-associated virus (AAV) with a double-floxed inverse open reading frame (DIO) was stereotactically injected unilaterally in substantia nigra of TH-cre mice to obtain cre-dependent expression of either HA-DAT-WT or HA-DAT-K619N selectively in TH-expressing dopaminergic neurons. As an additional control, we injected mice with an AAV construct encoding cre-dependent mCherry. Immunohistochemical stainings of midbrain- and striatal slices from TH-cre mice injected with AAV8-DIO-HA-DAT-WT, AAV8-DIO-HA-DAT-K619N, or AAV8-DIO-mCherry confirmed expression of the viral transgenes in TH-positive VTA neurons and revealed the presence of both HA-DAT-WT and HA-DAT-K619N in TH-labeled projections/axonal terminals in the striatum (Supplementary Figure 4), Interestingly, we observed that the effect of overexpressing HA-DAT-K619N differed markedly from HA-DAT-WT overexpression. First, mice expressing HA-DAT-WT displayed larger AMPH-induced hyperlocomotion compared to both HA-DAT-K619N and mCherry injected mice (Figure 3A, B). Secondly, the unilateral overexpression of HA-DAT-WT established rotational laterality with more contralateral rotations upon AMPH stimulation, whereas overexpression of HA-DAT-K619N did not establish AMPH-induced rotational laterality (Figure 3C-E). Note that in lesion studies, AMPH induces rotations toward the lesioned side (Bjorklund & Dunnett, 2019).

**Figure 4.**
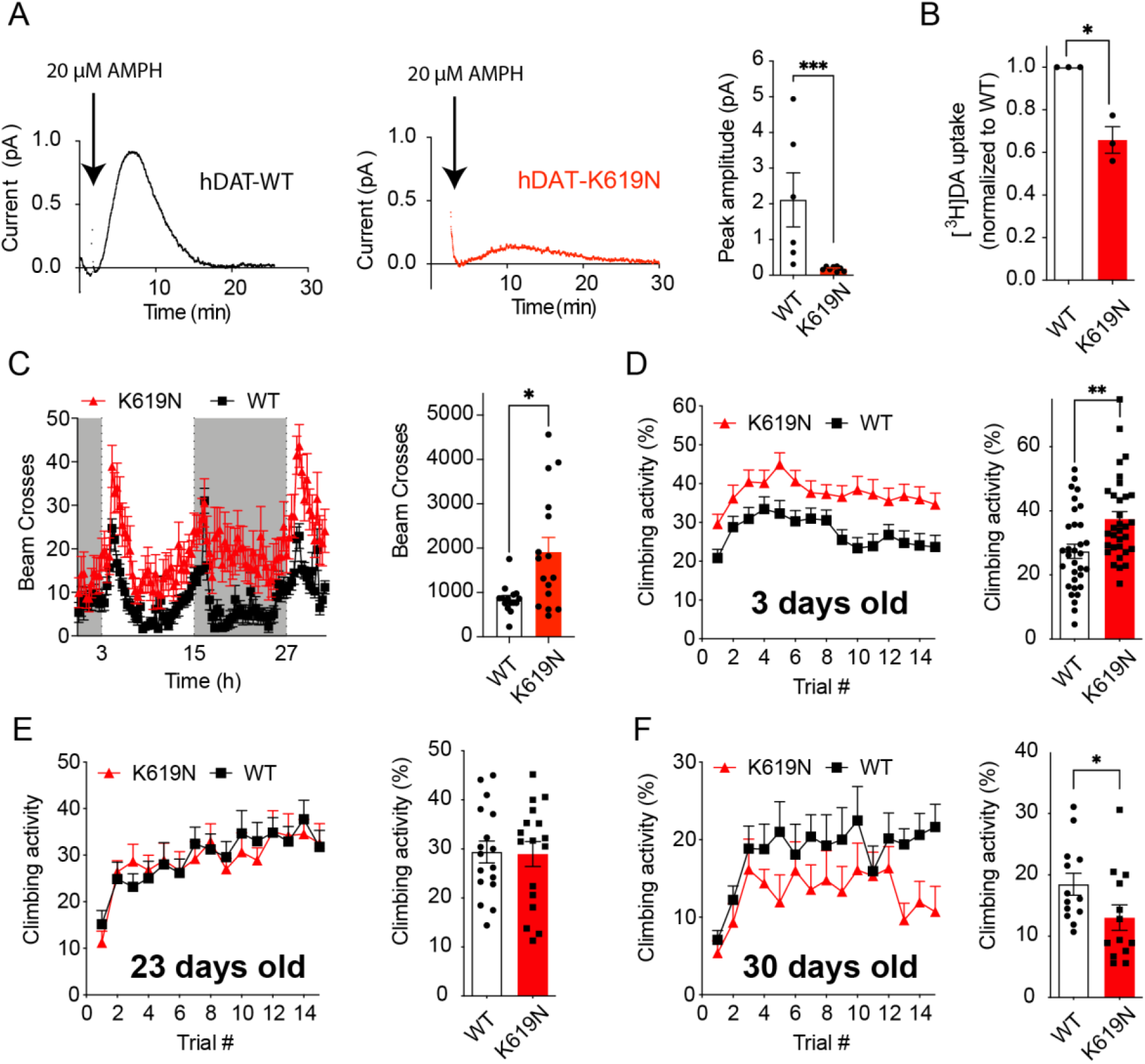
Expression of DAT-K619N drives dopaminergic dysfunction and progressive locomotor disturbances in *Drosophila*. (A) Amperometric recordings of AMPH-induced DA efflux in whole-brains from Drosophila expressing human DAT-WT or DAT-K619N. Representative traces and quantification of peak amperometric currents is shown. DAT-K619N flies display markedly reduced AMPH-induced efflux (p<0.001, Mann-Whitney test, N=6-8). (B) DA uptake (200nM for 15 min) into isolated whole fly brains, normalized to DAT-WT in each experiment. DA uptake is compromised in brains from the DAT-K619N strain compared to DAT-WT (p<0.05, one-sample t test, N=3).(C) Locomotor activity recordings of 3-5 days old flies over 30 h show that DAT-K619N flies are hyperactive during both light phase (light bars) and dark phases (dark bars), with a 125% increase in mean number of beam breaks (849±82 for DAT-WT vs 1910±320 for DAT-K619N, p<0.05, Mann-Whitney test, N=15-16 flies). (D-F) Assessment of negative geotactic crawling response in flies that are 3-5 days old (D), 23 days old (E) and 30 days old (F). The curves show climbing activity over 15 consecutive trials. Bar diagrams compare the mean climbing activity of the 15 trials. 3-5 days old DAT-K619N flies display a hyperactive phenotype (p<0.01 Mann-Whitney test, N=32 cohorts), which is absent at day 23 (p>0.05 Mann-Whitney test, N=17 cohorts) and at day 30, the DAT-K619N flies show reduced climbing activity compared to DAT-WT expressing flies (p<0.05 Mann-Whitney test, N=13 cohorts). This indicates that the DA dysfunction in DAT-K619N flies progresses over time.

### Expression of DAT-K619N drives dopaminergic dysfunction and progressive locomotor disturbances in Drosophila

Dopaminergic control of locomotion is highly conserved across species including *Drosophila* (Hamilton *et al.*, 2013; Pizzo *et al*, 2013). In another approach to investigate the impact of DAT-K619N *in vivo*, we used the Gal4/UAS system to generate knock-in (KI) lines, expressing a single copy of either EGFP-tagged or untagged human DAT-WT or DAT-K619N in dopaminergic neurons of *Drosophila* DAT (dDAT) KO flies. Confocal imaging of EGFP-tagged DAT-WT and DAT-K619N showed comparable expression of both transgenes in the somatic regions (PPL1) and in the fan-shaped body, which is a major projection target of these neurons (Supplementary Figure 5). Measurement of DA uptake in intact, isolated brains from the flies expressing untagged transporters, however, showed that uptake into DAT-K619N brains was reduced to 65.8±6% of DAT-WT brains (mean uptake DAT-WT: 333±5 fmol/brain vs 219±20 fmol/brain in DAT-K619N, Figure 4A). Moreover, in the DAT-K619N KI flies, AMPH-induced DA efflux was reduced by ~80% (DA peak current) compared to DAT-WT as determined by amperometric recordings (Figure 4B). Importantly, the apparent dopaminergic imbalance imposed by DAT-K619N also seemed sufficient to drive changes in locomotor activity, that is, when the flies were placed in activity chambers more beam breaks were recorded for DAT-K619N flies compared to DAT-WT flies during both the light and the dark cycle (Figure 4C). In addition, a test of startle-induced negative geotactic response demonstrated enhanced climbing activity for DAT-K619N (Figure 4D). To monitor if the locomotor dysfunction persisted or progressed with time, we evaluated also older flies. Remarkably, we found that the hyperactive climbing response of DAT-K619N expressing flies was lost in 24 days old flies, and by day 30 the hDAT-K619N flies were showing deficient climbing compared to DAT-WT expressing flies. This finding suggests a progressive nature of the dopaminergic dysfunction in the DAT-K619N flies (Figure 4D-F).

**Figure 5.**
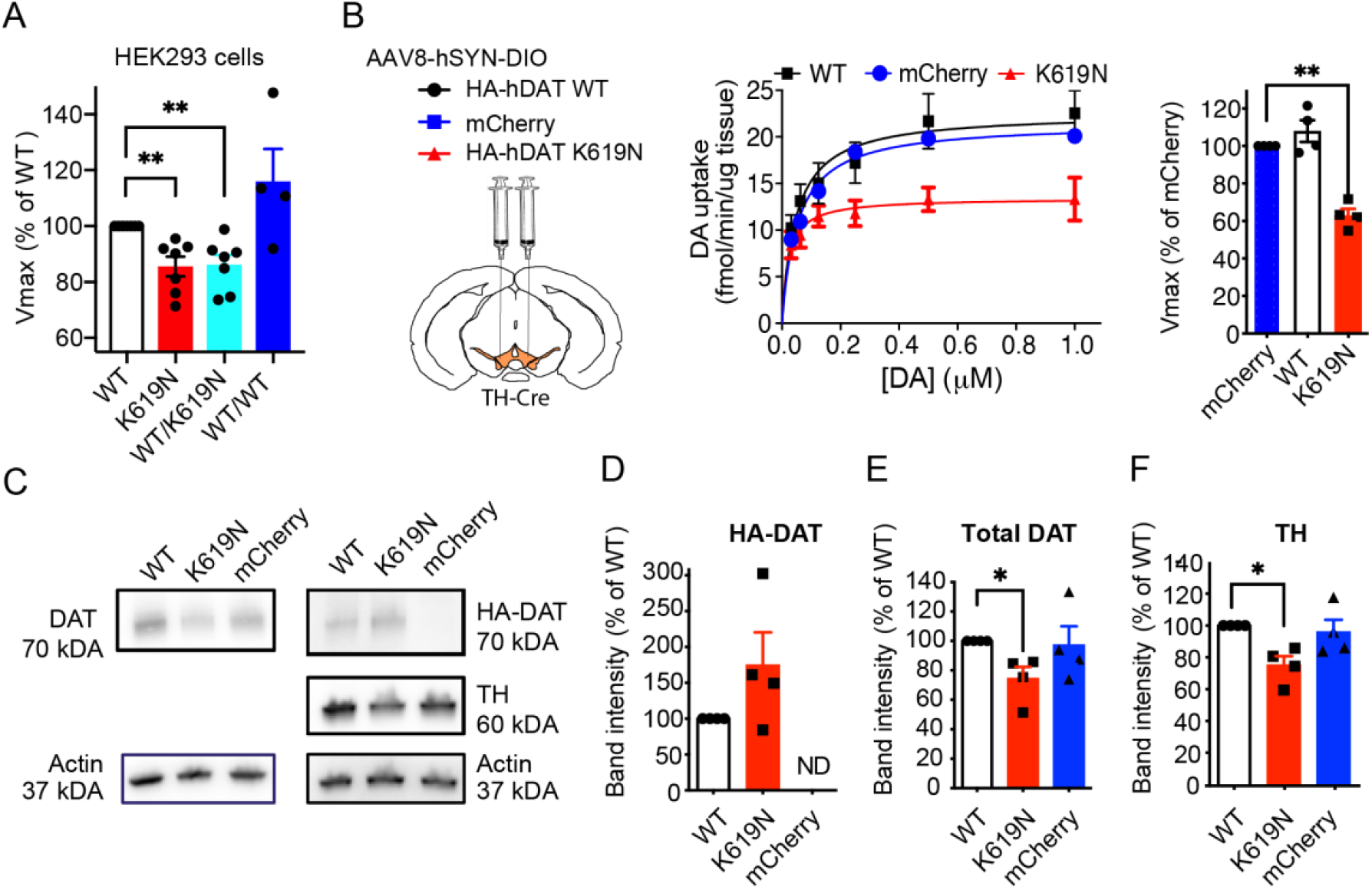
DAT-K619N exerts dominant-negative effect on DAT-WT. (A) Evaluation of dominant-negative effects of DAT-K619N *in vitro.* DA uptake was measured in HEK293 cells that were co-transfected with equal amounts (1.5μg) of DAT-K619N and DAT-WT, and compared to cells transfected only with DAT-WT (1.5μg 1.5μg empty vector), DAT-K619N (1.5μg 1.5μg empty vector), or with 3μg of DAT-WT as a control. Each experiment was performed in triplicates and normalized to V_max_ of DAT-WT (1.5μg 1.5μg empty vector). The V_max_ of DAT-K619N/DAT-WT co-transfected cells (1.5μg 1.5μg) is reduced relative to DAT-WT (1.5μg 1.5μg empty vector) to a level similar to that obtained for DAT-K619N alone (**p<0.01, one-sample t-test, N=4-7), indicating dominant-negative actions of DAT-K619N. (B) Evaluation of dominant-negative effects *in vivo* was done by selectively overexpressing HA-DAT-WT or HA-DAT-K619N in dopaminergic neurons, using bilateral midbrain AAV injections in TH-cre mice and performing [^3^H]-DA uptake on striatal synaptosomes. Injections with AAV encoding only mCherry was used to allow comparison with endogenous DA uptake levels. Uptake curves are average curves of four experiments each performed in triplicates. Quantification of V_max_ normalized to mCherry shows that HA-DAT-K619N reduces DA uptake below the endogenous uptake capacity, which is not seen for mice expressing HA-DAT-WT (p<0.01, one-sample t-test, N=4). (C) Western blotting analysis of the striatal synaptosomal preparations. Representative blots for total DAT, HA-DAT, and TH are shown. Membranes were stripped between blotting for HA-DAT and TH. Actin was detected to confirm equal loading. Specific detection of HA-DAT-WT and HA-DAT-K619N using an anti-HA antibody confirmed that both constructs are trafficked to striatal terminals. (D-F) Quantification of relative expression levels revealed a reduction in both total DAT expression (E) and TH (F) in HA-DAT-K619N-injected mice (*p<0.05, one-sample t-test, N=4), while HA-DAT did not show significant differences between HA-DAT-WT and HA-DAT-K619N-injected mice (P=0.21, one-sample t-test, N=4).

### K619N exerts dominant-negative impairments on DA uptake in vitro and in vivo

Given the heterozygote carrier status of the identified patients, an important question is whether a single DAT-K619N allele is sufficient to drive DA dysfunction and thereby potentially confer disease risk. DAT has been shown to form oligomers (Sitte *et al*, 2004), which could allow DAT-K619N to directly influence WT-DAT function or trafficking in heterozygote carriers. We therefore co-transfected HEK293 cells with equal amounts of DNA (1.5μg) of both DAT-K619N and DAT-WT and compared the resulting DA uptake capacity to that obtained when cells were transfected with only DAT-WT or DAT-K619N, using an empty vector to keep the total amount of DNA constant (3μg). Notably, the uptake capacity of co-transfected cells (1.5μg DAT-WT 1.5μg DAT-K619N) was significantly reduced compared to DAT-WT single transfection (1.5μg DAT-WT 1.5μg empty vector), and similar to that of cells transfected only with DAT-K619N (V_max_=86±3% of DAT-WT for WT/KN co-transfection and 87± 3% of DAT-WT for DAT-K619N alone). As a control, we transfected cells with 3μg of only DAT-WT, which did not establish a similar reduction in uptake (Vmax= 116 ±9 % of 1.5 μg WT), suggesting that the DAT-K619N variant has a dominant-negative effect on DAT-WT function (Figure 5A).

We sought next to evaluate the dominant-negative effect of the DAT-K619N variant *in vivo.* To do this, we expressed DAT-K619N in dopaminergic neurons of mice on top of the endogenously expressed DAT, thereby mimicking the heterozygote genotype of the patients. TH-cre mice received bilateral injections of AAVs encoding either HA-DAT-K619N, HA-DAT-WT or mCherry in VTA, and striatal synaptosomes were prepared to compare the resulting ^3^H-DA uptake capacity. Strikingly, mice expressing HA-DAT-K619N showed a pronounced reduction (~40%) in DA uptake relative to the endogenous uptake capacity derived from mice expressing the mCherry reporter, indicating a dominant negative effect of DAT-K619N *in vivo*. By contrast, mice injected with HA-DAT-WT did show significantly altered DA uptake relative to mice injected with mCherry (Figure 5B). The reduction in V_max_ observed in mice expressing HA-DAT-K619N was not accompanied by significant changes in K_M_ (K_M mCherry_=55±9nM, K_M K619N_=24±6 nM, and K_M DAT-WT_ =57±18 nM, p>0.05, one-way ANOVA, with Holm-Sidak posttest, N=4). By western blotting analysis of the synaptosomal fractions, we confirmed, by use of the HA-tag, the presence of both HA-DAT-WT and HA-K619N in the striatum (Figure 5C, D). Interestingly, quantification of the total DAT signal revealed a ~25% reduction in mice injected with HA-DAT-K619N compared to mice injected with HA-DAT-WT (Figure 5C, E). Moreover, we observed a reduction in TH expression in the synaptosomal fractions from HA-DAT-K619N expressing mice compared to HA-DAT-WT (75±6% of HA-DAT-WT, Figure 5C, F), further supporting that the HA-DAT-K619N variant generates changes in the DA system that are distinct from HA-DAT-WT.

### DAT-SPECT scans support progressive neurodegeneration

The apparent DA dysfunction imposed by hDAT-K619N in vivo prompted us to go back to patient 2 and compare DAT-SPECT scans acquired with an interval of seven years (2006 and 2013). The first scan demonstrated a clear reduction in DAT binding in striatum (caudate nucleus and putamen) with the largest reduction seen on the left hemisphere, consistent with a right-sided predominance of the patient’s symptoms. Although the scan resembles those observed in neurodegenerative parkinsonism (Tatsch & Poepperl, 2013), the loss of DAT availability could, in principle, reflect both dopaminergic cell loss and reduced DAT expression connected to the DAT-K619N variant. However, the second scan acquired seven years later, showed an accelerated loss of [^123^I]-FP-CIT binding, compared to the expected decline from age-matched controls (Figure 6). This suggests that loss of DA neurons is contributing to the loss of DAT binding, and thus that dopaminergic neurodegeneration is part of the neuropathology in patient 2.

**Figure 6.**
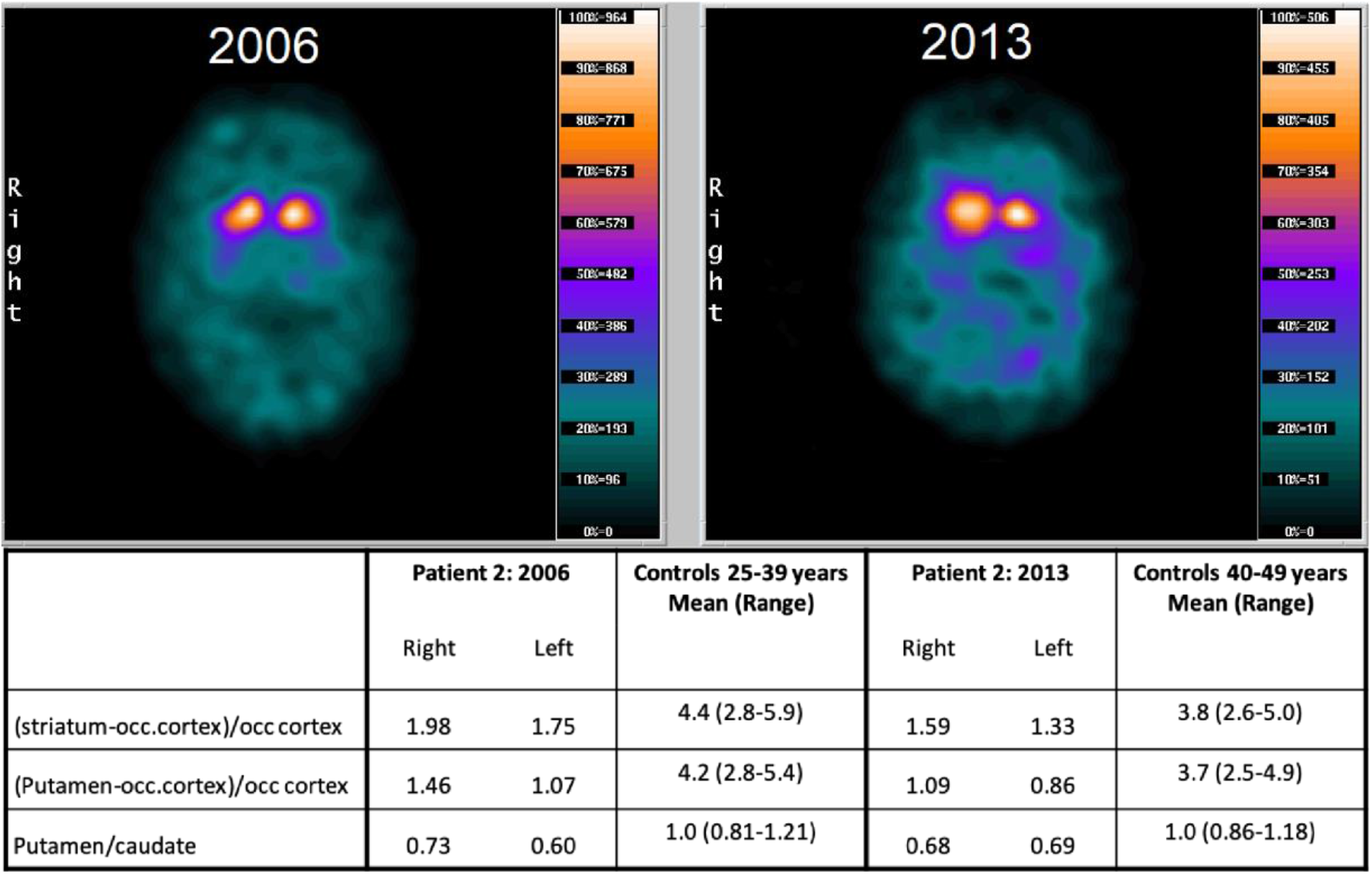
[^123^I]FP-CIT SPECT imaging show progressive loss of DAT binding. (A) [^123^I]FP-CIT SPECT imaging of patient 2 carrying the DAT-K619N variant. DAT scans of patient 2, acquired seven years apart, suggest progressive neurodegeneration. Images were taken with identical procedures on the same scanner at age 34 (in 2006, left) and age 43 (in 2013, right). DAT availability was quantified as the ratio between specifically bound radioligand and non-specific radioligand binding in occipital cortex. The following uptake ratios were calculated: (1) specific uptake in entire striatum= (striatum-occipital cortex)/occipital cortex, (2) Specific uptake in putameN= (putamen-occipital cortex) /occipital cortex, and (3) putamen/caudate nucleus to evaluate symmetry/asymmetry of DAT availability.

## Discussion

Diseases involving DA dysfunction span both neurological disorders such as parkinsonism and widespread neuropsychiatric diseases such as ADHD, ASD, bipolar disorder, schizophrenia, and depression (Cousins *et al.*, 2009; Del Campo *et al.*, 2011; Dichter *et al.*, 2012; Howes & Kapur, 2009; Iversen & Iversen, 2007; Robison *et al.*, 2020). DAT is a powerful regulator of DA neurotransmission, but the genetic and mechanistic link between insult to DAT function and DA-related pathologies is not clear. Specifically, the role of heterogeneity for rare coding variants in parkinsonism and/or neuropsychiatric disease remains elusive. Here, we provide new insight into the link between DAT dysfunction and human disease by employing a translational approach to investigate the rare coding variant DAT-K619N, which we identify in an index patient with early-onset parkinsonism as well as in multiple patients with psychiatric disorders. We combine detailed clinical data, neuroimaging, and large-scale exome data to address the occurrence and phenotypic spectrum of DAT-K619N carriers, and employ *in vitro* studies alongside *in vivo* and *ex vivo* investigations in mouse and *Drosophila* models of DAT-K619N to uncover the functional consequences of the rare, but recurring coding variant with apparent dominant-negative properties.

Our in vitro studies in heterologous cells revealed a parallel reduction in DA uptake, amphetamine-induced DA efflux, and [^3^H]-CFT binding capacity for DAT-K619N compared to DAT-WT, which, together with preserved Zn^2+^-dependent regulation and normal ligand and ion affinities, suggested that the K619N mutation changes the cellular DAT processing without imposing conformational alterations that impair catalytic activity. The phenotype was supported by whole-cell surface biotinylation experiments and surface staining of DAT in live cells, which demonstrated a decrease in surface-expressed DAT-K619N compared to DAT-WT. Moreover, live cell imaging of DAT-K619N coupled to a fluorescent timer showed that the DAT-K619N protein, on average, was younger (smaller red-to-blue ratio) than DAT-WT, and we also observed increased localization of mCherry-tagged DAT-K619N to lysosomes in live CAD cells, suggesting that DAT-K619N is subject to accelerated turnover leading to lower expression of active transporter. Interestingly, the DAT C-terminus, including the C-terminal PDZ-binding motif that K619 is part of (L618, K619 and V620), has been subject to several previous mutational studies (Bjerggaard *et al.*, 2004; Miranda *et al*, 2004; Rickhag *et al.*, 2013; Torres *et al*, 2003; Torres *et al.*, 2001). We previously alanine-substituted, the PDZ-binding sequence (DAT-AAA), which, conceivably due to disruption of interactions with yet unknown PDZ domain scaffold proteins, led to reduced expression and enhanced degradation of the transporter, and, hence, a mutant phenotype similar to what we find for DAT-K619N (Rickhag *et al.*, 2013). Other mutations have implicated the same PDZ-binding sequence in folding and ER export (Bjerggaard *et al.*, 2004; Torres *et al.*, 2001), collectively supporting that the DAT C-terminus is highly sensitive to mutational changes and that an intact C-terminus is critical for biosynthesis, trafficking and turnover of the DAT protein.

Our subsequent *in vivo* and *ex vivo* experiments supported that the DAT-K619N variant can give rise to changes in DA function of potential pathological importance. The experiments also supported that the molecular phenotype is markedly more pronounced *in vivo*, consistent with previous finding for the DAT-AAA mutation (Rickhag *et al.*, 2013). Thus, we found that whole-brain DA uptake was reduced by ~35% and amphetamine-induced DA efflux was reduced by more than 80% in flies expressing the human DAT-K619N variant compared to DAT-WT. Moreover, young flies expressing DAT-K619N demonstrated hyperlocomotion and an enhanced negative geotactic response compared to DAT-WT-expressing flies. In mice, the differential effect of DAT-K619N and DAT-WT was sufficient to establish differences in DA-directed behaviors, which we as demonstrated by assessing AMPH-induced rotations in mice with unilateral overexpression of HA-DAT-WT or HA-DAT-K619N in nigrostriatal DA neurons. Rotational behavior is sensitive measure of asymmetry in striatal DA (Bay Konig *et al.*, 2019; Bjorklund & Dunnett, 2019), and we observed that while unilateral HA-DAT-WT expression led to both increased overall locomotor activity and contralateral rotational bias in response to AMPH, this was not observed in mice expressing HA-DAT-K619N or the mCherry control. Yet another important finding was the ~40% reduction in striatal synaptosomal uptake seen upon bilateral viral expression of HA-DAT-K619N, but not upon expression of HA-DAT-WT or mCherry in midbrain DA neurons. This suggests that DAT-K619N is capable of inhibiting DA uptake in a dominant-negative manner, as was observed *in vitro* too (Figure 5). From a mechanistic perspective, such a dominant-negative effect of DAT-K619N is likely linked to transporter oligomerization (Hastrup *et al*, 2001; Torres *et al.*, 2003) meaning that accelerated degradation of DAT-K619N through interaction with DAT-WT might lead to concomitant accelerated degradation of DAT-WT. Notably, it was previously found that non-functional C-terminal truncations in DAT can reduce surface expression of the DAT-WT in a dominant-negative fashion, and it was concluded that this was the result of oligomeric interactions between the mutants and the WT protein (Torres *et al.*, 2003).

The dominant negative action of DAT-K619N is particularly noteworthy because patient 2, to our knowledge, is the first example of a heterozygote carrier of a coding DAT variant with early-onset parkinsonism. To date, all previously described patients with DTDS have been either homozygous or compound heterozygote for disruptive variants in *SLC6A3* (Hansen *et al.*, 2014; Heidari *et al*, 2020; Kurian *et al.*, 2011; Kurian *et al.*, 2009; Ng *et al.*, 2014; Yildiz *et al*, 2017). Thus, the current data might conceptually and mechanistically expand the allelic disease spectrum of DTDS from biallelic non-synonymous variants to single susceptibility/disease alleles. It is, however, not clear exactly how much DAT activity is required to maintain human health. In the classical form of DTDS, where biallelic mutations cause complete loss of DAT function, the disease manifests already in early infancy (Kurian, 1993). On the other hand, we recently described an atypical case of DTDS where the patient was compound heterozygote for partially disruptive DAT variants with an estimated residual DAT activity of ~30% (Hansen *et al.*, 2014). This patient, like patient 2 in the present study, suffered from early-onset parkinsonism and neuropsychiatric disease and was classified as suffering from atypical DTDS. Also, three brothers with atypical juvenile-onset DTDS have been described for which the predicted DAT activity was ~8% (Kurian, 1993; Ng *et al.*, 2014), strongly supporting a phenotypic spectrum determined by residual uptake activity of DAT. The final level of DAT uptake capacity in human heterozygote carriers of DAT-K619N is difficult to predict and may vary between people and be influenced by unknown gene-gene or gene-environment interactions. Our functional studies strongly suggest that heterozygosity for DAT-K619N would considerably compromise, although not completely abolish, DA uptake, furthermore supported by the DAT-SPECT scans of our index patient.

Another question is whether impairments in DAT function can contribute to a neurodegenerative process. The SPECT scans performed seven years apart showed accelerated loss of DAT binding, compared to age-matched controls, indicative of progressive dopaminergic neurodegeneration. This finding aligns with previous data from our other patient with atypical, adult-onset DTDS, in whom we also found evidence for dopaminergic degeneration (Hansen *et al.*, 2014). Interestingly, when HA-DAT-K619N was expressed in WT mice, we observed not only a decrease in uptake but also a parallel reduction in DAT and TH levels, as assessed by immunoblotting. Reduction in TH is often considered indicative of dopaminergic denervation or neurodegeneration, which is further strengthened by the parallel reduction in DAT (Luk *et al*, 2012; Masliah *et al*, 2000; Matheoud *et al*, 2019; Steinkellner *et al*, 2018). It is possible, however, that the reduced TH level instead is compensatory to reduced DAT function and elevated extracellular DA. We should also note that when expressing HA-DAT-K619N unilaterally in nigrostriatal neurons we did not observe AMPH-induced ipsilateral rotational bias, an often-used measure of motor impairments and functional recovery following unilateral lesions. At the same time, it is important to point out that this phenotype is an indirect correlate for dopaminergic neurodegeneration and it only manifests when ~50% or more of nigral DA neurons are lost (Bjorklund & Dunnett, 2019). Of further interest, the behavioral data on flies expressing DAT-K619N suggest that the dopaminergic dysfunction caused by this mutant is not static but rather progressive in nature. In the hDAT-K619N flies, we observed an initial hyperactive climbing response, which disappeared over time, and at day 30, the climbing activity had even dropped below that of WT hDAT KI flies (Figure 4). Further studies are needed to elucidate if the loss of DA neurons that have now been observed in two independent cases of atypical DTDS reflects a shared pathophysiological mechanism. Our findings provide additional support to the notion that DAT dysfunction may be directly involved either in initiating neurodegenerative processes in DA neurons or in aggravating the selective vulnerability of these highly specialized neurons.

The genetic component of neuropsychiatric diseases has been firmly established (Gratten *et al*, 2014) and shown to involve both common variants and rare variants (Bray & O’Donovan, 2019; Gratten *et al.*, 2014; Sullivan *et al*, 2012). Identification of causal variants and interrogation of their functional impact are critical next steps for further advancing our disease understanding. *SLC6A3* has not been identified as a risk gene in GWAS studies of psychiatric disease or neurological disorders (MacArthur J, 2017). This, however, does not infer that DAT dysfunction cannot generate disease-causing, contributing, or modulating processes. Rather, it reflects the absence of common risk variants. Several rare coding DAT variants with functional deficits both *in vitro* and *in vivo* have been identified in patients with psychiatric disorders (Bowton *et al.*, 2014; Campbell *et al.*, 2019; Cartier *et al.*, 2015; Davis *et al*, 2018; DiCarlo *et al.*, 2019; Gowrishankar *et al*, 2018; Grunhage *et al.*, 2000; Hamilton *et al.*, 2013; Hansen *et al.*, 2014; Herborg *et al.*, 2018; Mazei-Robison *et al.*, 2005; Sakrikar *et al.*, 2012; Stewart *et al*, 2019; Wu *et al.*, 2015) but the causal link between these putative risk alleles and disease is still elusive as the family trees and cohort sizes have been too small for meaningful linkage or association analysis. We have, to the best or our knowledge, carried out the first association analysis of a coding DAT variant using large-scale exome data, and demonstrate a nominally significant association to bipolar disease. Of notice, we found that the estimated odds ratios for DAT-K619N were increased across all the diagnostic categories and also for the cross-disease association analysis, which could imply that DAT-K619N, and possibly other disruptive mutations can act as shared risk factors for neuropsychiatric disease. The identification of DAT-K619N carriers among both healthy subjects and patients with different neuropsychiatric diseases is not surprising. Pleiotropic effects and incomplete penetrance have been repeatedly described for both common and rare variants in psychiatric disorder even for variants with OR as high as 60. It, therefore, seems likely that the phenotypic outcome of DAT-K619N is dependent on other factors, such as genetic background and environmental factors (Bray & O’Donovan, 2019). Larger samples are needed to establish true associations and to more accurately determine the relative penetrance of BP, schizophrenia, ADHD, ASD and depression. In addition, the study population is relatively young and it would be relevant with follow-up studies to uncover re-diagnoses, potential new incidences of disease among control subjects and investigations of movement disorder later in life.

In summary, our investigations identify a novel rare coding DAT variant in the critical C-terminal PDZ-binding motif that even when present on a single allele can cause dopaminergic dysfunction through a dominant-negative effect and thereby conceivably confer risk for both neuropsychiatric disease and neurodegenerative early-onset parkinsonism. This insight should be crucial in future efforts aimed at further understanding how altered DAT function contribute to DA pathologies and how we can develop better treatment strategies for these diseases.

## Materials and methods

### Subjects: clinical assessment and procedures

Patient 1 was part of the Simon Simplex Collection which comprises sequencing data from more than 2000 ASD families as previously described (Fischbach & Lord, 2010). Patient 2 was identified in a previously described referral-based hospital cohort of patients with early-onset parkinsonism or related atypical movement disorders (Hansen *et al.*, 2014). Medical case notes were reviewed to delineate the clinical history and prior treatments. Video recordings were made for documentation of clinical features. A DAT-SPECT scan was made for comparison with a DAT-SPECT scan made seven years prior. The two scans were performed on the same scanner, meticulously in the same manner. The patient received 200 mg sodium perchlorate i.v. to block uptake of free ^123^I in the thyroid gland before i.v. injection of 185 MBq [^123^I]-2ß-carbometoxy-3ß-[4-iodophenyl]-N-[3-fluoropropyl]nortropane ([^123^I]FP-CIT, GE Healthcare, Amersham, UK). The SPECT data acquisition was begun after exactly 3 hours, with a PRISM 3000XP (Marconi, Phillips) triple-headed gamma camera equipped with low-energy, ultra-high-resolution fan-beam collimators. Image reconstruction was performed using iterative reconstruction with scatter and non-uniform attenuation correction. Pixel size after reconstruction was x,y=3.1×3.1 mm^2^ with a slice thickness of 6.2 mm. From the individual reformatted dataset, the two neighboring striatal slices with maximum counts per pixel were summed and used for image analysis. DAT availability was quantified as the ratio between specifically bound radioligand and non-displaceable radioligand for five regions of interest (ROIs) representing striatum (caudate nucleus and putamen) bilaterally and in the occipital cortex for calculation of non-specific binding. The following uptake ratios were calculated: (1) specific uptake in striatum= (striatum-occipital cortex)/occipital cortex, (2) specific uptake in putamen =(putamen-occipital cortex) /occipital cortex, and (3) putamen/caudate nucleus to evaluate symmetry/asymmetry of DAT availability.

### iPSYCH exome sequencing data

The exome sequencing data used in the present study are from the integrated psychiatric research (iPSYCH) consortium’s first phase genotyping of a nation-wide Danish birth cohort of 1,472,762 individuals born between May 1, 1981, and December 31, 2005. The iPSYCH case-cohort design has been described in greater detail previously (Pedersen *et al.*, 2018). The study was approved by the Danish Data Protection Agency. Information about the occurrence of DAT-K619N among cases and controls rely on whole exome sequencing data from a subset of 19,851 samples from the first phase of iPSYCH case-cohort study. Procedures for exome sequencing, sample and variant quality control are described in Supplementary Methods. For carrier-based association analysis, individuals within the case and the control samples were categorized as either carriers or non-carriers and 2×2 contingency tables with two-sided Fisher’s exact tests were applied to test for carrier-based disease associations. A total of 6 association tests was performed and study-wide significance is accordingly adjusted to P<0.0083.

### Cell culturing and transfection

Human embryonic kidney cells, Cath.-a-differentiated (CAD) cells, and Cos-7 cells were maintained and transfected as previously described (Hansen *et al.*, 2014; Herborg *et al.*, 2018). Assays were conducted 36-48 h after transfection. All *in vitro* experiments, except live cell imaging of fluorescently-tagged constructs, were carried out on heterologous cells transiently transfected with pRC/CMV-hDAT-WT or hDAT-K619N. Fluorescently-tagged constructs (mCherry and slowFT) were generated by N-terminal fusion of the fluorescent protein to a synthetic hDAT in pcDNA3. The slow fluorescent timer was kindly provided by Dr. Verkhusha (Albert Einstein College of Medicine, New York, USA). The K619N mutation was introduced in the various hDAT background constructs using either QuickChange site-directed mutagenesis (Stratagene) or two-step PCR. All constructs were verified by sequencing.

### [^3^H]-DA uptake and [^3^H]-CFT binding experiments in vitro

[^3^H]-DA uptake experiments for determination of kinetic constants, ligand affinities, and ion- and Zn^2+^-dependent regulation, as well as [^3^H]-CFT binding experiments, were carried out as described in (Herborg *et al.*, 2018). The functional characterization of DAT-K619N in Cos-7 cells was carried out alongside a head to head comparison of previously reported disease-associated coding DAT variants published in (Herborg *et al.*, 2018) and DAT-WT data is therefore presented as a dotted line. To evaluate dominant-negative effects of the DAT-K619N variant on DAT-WT function, [^3^H]-DA uptake experiments were performed on HEK293 cells, that were co-transfected with DAT-K619N and DAT-WT (1.5 μg of each construct), and compared with cells that were transfected with only DAT-K619N or DAT-WT (1.5 μg 1.5 μg empty DNA) as well as with cells transfected with 3 μg of DAT-WT as a control. Thus, the total DNA amount was kept constant at 3 μg for all conditions. Michaelis-Menten kinetics was used to fit saturation uptake data. Competition uptake and binding experiments were fitted by non-linear regression with variable slope. Ion dependency uptake curve were fitted by one-site specific binding model.

### In vitro amperometry recordings

AMPH-induced efflux was assessed by amperometry recordings on transiently transfected HEK293 cells expressing DAT-WT or DAT-K619N as described previously (Hansen *et al.*, 2014).

### Live confocal microscopy of heterologous cells

Live imaging of transiently transfected HEK293 and CAD cells was done in imaging buffer (25 mM HEPES, pH 7.4, with 130 mM NaCl, 5.4 mM KCl, 1.2 mM CaCl2, 1.2 mM MgSO4, 1 mM L-ascorbic acid, 5 mM D-glucose) and cells were reversely transfected in 8 well Lab-Tek II. Confocal images were obtained either on a Zeiss LSM 510 or a Zeiss LSM 780 point-scanning confocal microscope (Zeiss) using an oil immersion 63×1.4 numerical aperture objective. Visualization of fluorophores and fluorescent proteins was achieved using the following laser/filter combinations:JHC 1–64 and mCherry: 543-nm helium-neon laser/ 560-nm long-pass filter; LysoTracker fluorophore and MFZ-9-18: 488 nm argon laser/494-565 nm bandpass filter; blue form of SlowFT: 405 nm diode laser/ 410-480 nm bandpass filter; red form of SlowFT: 580 nm in-tune laser/ 582-689 nm bandpass filter. Confocal images for all experimental series were acquired from at least three independent transfections.

For live cell imaging with JHC 1-64 (Eriksen *et al.*, 2009; Hansen *et al.*, 2014), cells were incubated 20 min with 20nM JHC 1-64 at room temperature in imaging buffer and washed three times before imaging. Quantification of the mean JHC 1-64 signal was done in ImageJ. The automated ‘Li’ threshold function was applied, and mean intensity was measured. The mean intensity of all images was normalized to the WT average mean intensity for each experiment. Spatiotemporal visualization of DAT-WT and DAT-K619N using N-terminal tagging with SlowFT was done in CAD cells. The blue-to-red SlowFT is characterized (*in vitro)* by chromophore maturation rates of 9.8 h for maximal blue fluorescence and 28h for half-maxima of the red fluorescence (Subach *et al.*, 2009). As the blue form of the timer is sensitive to photoconversion (Subach *et al.*, 2009), we used a green fluorescent cocaine analogue, MFZ-9-18 to label surface DAT (WT or K619N) and to allow identification of transfected cells without direct imaging of the timer. Thus, cells were washed in imaging buffer and incubated 20 min with 400 nM MFZ-9-18 (RT) and washed twice before imaging. Transfected cells were identified using the MFZ-9-18 signal, and images of the red form of the slowFT were acquired first to minimize confounding effect of blue-to-red photoconversion (Subach *et al.*, 2009). Images were analyzed in the Fiji software. Identical settings were used to threshold all images of SlowFT-DAT-WT and SlowFT-DAT-K619N within a given imaging session. Pixels with values below the threshold were assigned with ‘NaN’ (not a number value), thereby excluding them from analysis. The red-to-blue ratio for each pixel was calculated. Next, we used the MFZ-9-18 image to generate a mask to isolate the ‘surface’ DAT fraction, and this picture was then subtracted from the total DAT red-to-blue ratio image in the Image Calculator function to isolate the intracellular fraction. Finally, to exclude pixels with red-to-blue ratio of zero a threshold value of 1*10^−10^ was applied, and the mean red-to-blue ratio was measured for the total-, surface-, -and intracellular DAT fractions and normalized to the WT average for each individual experiment.

Imaging of mCherry-DAT-WT or mCherry-DAT-K619N was done in live CAD cells. Images were acquired of both mCherry alone and following 15 min incubation LysoTracker^®^ Green DND-26 (ThermoFisher Scientific) to investigate lysosomal targeting. The mean intensity of the mCherry signal was quantified as described for JHC 1-64. The fractional overlap between mCherry and LysoTracker positive compartments (Manders coefficient) was calculated using the JaCoP Plug-in for ImageJ.

### Mice

TH-Cre mice were obtained from Jackson Laboratory (stock number: JAX:8601, strain name: B6.Cg-Tg(TH-Cre)1Tmd/J) and backcrossed with C57BL/6N mice for at least seven generations, and maintained in a hemizygous state. All experiments were performed on adult female mice in accordance with guidelines from the Danish Animal Experimentation Inspectorate (#2017-15-0201-01160)

### AAV constructs and stereotactic surgery

Selective expression of human DAT-WT or DAT-K619N in dopaminergic neurons *in vivo* was achieved by stereotactic injections of cre-dependent AAV8 constructs into the midbrain of adult female TH-Cre mice (10-14 weeks). To allow detection of only the virally encoded DAT-WT and DAT-K619N, an HA-tag was introduced in the second extracellular loop (Sorkina *et al*, 2006). The viral vectors: AAV8-hSyn-DIO-HA-hDAT-WT, AAV8-hSYN-DIO-HA-hDAT-K619N, were cloned in-house and manufactured from Vector Biolabs. The AAV8-hSYN-DIO-mCherry vector was purchased from Addgene (plasmid #50459). All three viral vectors had egual titers of 4.0×10^12 vg/ml.

TH-Cre mice were deeply anaesthetized using induction flow of 2% isoflurane and 0.5% oxygen and placed in stereotaxic head holder (Kopf instruments) with fitted anesthesia mask. Anaestesia was maintained using 1-1.5% isoflurane in 1% oxygen. The following bregma coordinates (in mm) were used to deliver unilateral or bilateral injections of 300-500nL of AAV into the midbrain: VTA: AP-3.3, ML±0.5,DV-4.5 (300nL); Substantia nigra dual injections: AP-3.0, ML±1.2, DV-4.5 (200nL) and AP-3.16, ML±1.6, DV-4.5 300nL).

### Synaptosomal uptake

DA uptake experiments on crude striatal synaptosomes from TH-cre mice bilaterally injected in VTA with AAV8-hSYN-DIO-HA-hDAT-WT, AAV8-hSYN-DIO-HA-hDAT-K619N, or AAV8-hSYN-DIO-mCherry was performed 4-5 weeks after injections as described in (Jensen *et al*, 2017). DA uptake kinetics was analyzed using a two-fold dilution row (1-0.031 μM) containing a mixture of unlabeled DA and 2, 5, 6-[^3^H]-DA (Perkin Elmer Life Sciences, USA) as previously described (Jensen *et al.*, 2017). Data was fitted with Michaelis-Menten.

### Immunohistochemistry on mice

Mice were transcardially perfused with 4% paraformaldehyde in 0.1 M PBS (pH 7.4) 5-6 weeks after viral injections. Coronal sections (40μm) from striatum and midbrain were rinsed in PBS and incubated 30 min in retrieval buffer (10mM trisodium citrate in water, pH 6.0) at 80 °C. After washing, sections were blocked for 30 min (PBS with 5% goat serum, 1% bovine serum albumin, 0.3% Triton X-100 in PBS) and incubated overnight (4 °C) with primary antibodies against TH (OPA1-04050 1:1000, Thermo Scientific) and HA (3F10 1:200, Roche). Sections were rinsed three times in washing buffer (0.25% bovine serum albumin and 0.1% Triton X-100 in PBS) and incubated 60 min with secondary Alexa488 and Alexa568 antibodies (ab150077 and ab175476, Abcam, 1:400 in washing buffer). Sections from mice injected with AAV8-hSYN-DIO-mCherry were stained only for TH. Sections were mounted on glass coverslips (Mentzel-Gläzer, 24×60 mm, Germany) using ProLong™ Gold Antifade Mountant mounting media with DAPI (P36931, Life Technologies). Images were acquired using a Slide Scanner Axio Scan.Z1 (Zeiss) with 488 nm, 561 nm, and DAPI channel settings.

### Immunoblotting

Whole cell lysates from transiently transfected HEK293 cells and crude striatal synaptosomes were made as previously described (Hansen *et al.*, 2014; Rickhag *et al.*, 2013). After adjustments of protein concentrations, equal amounts of protein were incubated with loading buffer (100mM DTT, 4x SDS loading buffer) for 1 h at 37°C. Samples were then resolved by SDS-PAGE and western blotting was carried out as described in (Hansen *et al.*, 2014) using the following antibodies: DAT (rat MAB369, 1:1000, Millipore) or HA (rat 3F10, 1:200, Roche). Following detection of HA-DAT in synaptosomal samples, the membranes were stripped in stripping buffer (62.5 mM Tris-HCl, 2% SDS, 2.5% dithiothreitol (DTT)) at 50°C for 30 min, blocked again, and re-probed for TH using rabbit anti-TH (OPA1-04050, 1:1000, Thermo Scientific). An HRP-conjugated anti-β-actin antibody (1:40.000, Sigma) was used as loading control. Band intensities were quantified using Fiji software.

### AMPH-induced rotations

TH-cre mice were stereotactically operated and received two AAV injections unilaterally (left) in lateral VTA and substantia nigra with either AAV8-hSYN-DIO-HA-hDAT-WT AAV8-hSYN-DIO-HA-hDAT-K619N, or AAV8-hSYN-DIO-mCherry. Three weeks later, the mice were placed in open-field chambers for 1.5 h of baseline locomotor recordings, after which they were given an intraperitoneal injection of 5mg/kg AMPH and placed back in the chamber for another 1.5h of recording. Videos were analyzed in Ethovision to obtain baseline and amphetamine-induced locomotion and ipsi/contralateral rotations. Experiments and data analysis were carried out in a ‘blind’ manner with the investigator unaware of the AAV manipulation.

### Drosophila genetics

*Drosophila* strains were reared and maintained on either standard cornmeal-molasses media or on Nutri-Fly™ Bloomington Formulation medium (Genesee Scientific, San Diego, California) at 25°C and under a 12:12 h light-dark schedule. Transgenic flies expressing untagged versions of human DAT-WT and DAT-K619N were used for behavioral testing, ^3^H-DA uptake and amperometry and generated, using the pBI-UASC vector (Wang *et al*, 2012) as described in (Hamilton *et al.*, 2013). *DAT*^*fmn*^ (dDAT KO), *TH-Gal4*, and *UAS-mCherry* lines were outcrossed to a *w*^*1118*^ control line for 10 generations and selected by PCR or eye color. Likewise, flies containing the untagged UAS-DAT-WT and UAS-DAT-K619N transgenes were outcrossed to *DAT*^*fmn*^ flies (in a *w*^*1118*^ background) for 10 generations before use and maintained in this manner. Age-paired adult male flies were used for all subsequent experiments. For visualization of the DAT-WT and DAT-K619N transgenes pUASTattB-GFP-hDAT-WT and pUASTattB-GFP-hDAT-K619N constructs were generated by GenScript (Nanjing, China). The transgenes were inserted into the P[CaryP]attP2 site on the third chromosome using Phi31C transformation to ensure equal levels of expression. Embryo injections and selection of transformats were performed by BestGene Inc. Other fly stocks include *w*^*1118*^ ((Bloomington Indiana Stock Center (BI) 6326), TH-GAL4 (BI 8848), UAS-mCherry-CAAX (Kyoto Stock Center 109594), and M[vas-int.Dm]ZH-2A; (M[3xP3-RFP.attP’]ZH-22A (Bl 24481) and *DAT*^*fmn*^ (dDAT KO). Genotypes of the knock-in flies used for imaging of GFP-hDAT in dopaminergic neurons in fly brains were (WT: *w; DAT*^*fmn*^; *TH-Gal4, UAS-mCherry-CAAX/UAS-GFP-hDAT-WT*) and (K619N: *w; DAT*^*fmn*^; *TH-Gal4, UAS-mCherry-CAAX/UAS-GFP-hDAT-K619N*). Both male and female flies were used in imaging experiments, but were subsequently separated in the analysis.

### Drosophila immunostaining, imaging and analysis

Brains from 2-5 days old adult flies were dissected in PBS and briefly stored in Schneider’s insect cell medium (Life Technologies, A820) supplemented with 5% heat-inactivated FBS prior to fixation in 4% paraformaldehyde in PBS for 45 min at RT. Following 6× 10 min washes in PBS with 1.0% Triton X-100 (PBX), the brains were incubated in blocking solution (PBX with 3% BSA and 2% goat serum) for 2 h (RT) and then with an anti-GFP VHH single domain antibody/nanobody (Chromotech, gt-250) custom-conjugated to Alexa 647 (1:200) in blocking solution for ~72 h at 4°C. Specimens were then washed 10x 10 min in PBX and fixed again in 3.7% formaldehyde in PBS for 45 min at RT, followed by two rinses and two 5 min incubations in quenching solution consisting of 20 mM Glycine and 50 mM NH_4_Cl in PBS. After a 5 min wash in PBS, brains were mounted in ProLong® Gold antifade reagent (Life Technologies, P36934) with the posterior side towards the coverslip.

Fly brains were imaged on a LSM700 confocal microscope using a Plan-Apochromat 20x/0.8 NA air objective. Anti-GFP-conjugated Alexa 647 was excited using a 639 nm diode laser and detected using a 640 nm long-pass filter. Fixed mCherry was detected directly without the use of antibodies using a 555 nm diode laser for excitation and a 578-700 nm detection window in a separate track.

The immunosignal from GFP-hDAT/GFP-hDAT^K619N^ and the fluorescent signal from mCD8-mCherry were quantified in confocal stacks using Fiji/ImageJ. Sum projections of mCD8-mCherry stacks were subjected to rolling ball background subtraction, thresholded using the 88^th^ percentile and a region of interest (ROI) was drawn manually to surround the fan-shaped body. Particle analysis was then applied to refine the ROI to relevant anatomic structures, and the integrated signal from anti-GFP and mCherry was measured.

### Drosophila behavioral testing

Basal activity measurements were made on 3 days posteclosion male flies, which were placed in tubes with food for 30h. Activity was measured by beam breaks and analyzed using Trikinetic software.

Startle-induced negative geotactic responses were assessed in cohorts of 3 days, 24 days, and 31 days old DAT-WT and DAT-K619N flies, as described in (Barone & Bohmann, 2013). Briefly, cohorts (~13-15 flies pr. tube) of DAT-WT and DAT-K619N flies were transferred to 15 cm cylinder glass tubes and habituated for 15 min. Negative geotactic responses were induced by three taps and climbing activity was derived as the percentage of flies passing a 6 cm line after 5 sec in young (3 days) flies, or after 10 sec in older flies. Each cohort was tested in 15 trials with a one-minute inter-trial period. At least 13 cohorts of each genotype were assessed at any time point and the mean climbing activity was compared for DAT-WT and DAT-K619N.

### Drosophila ^3^H-DA uptake and amperometry assays

DA uptake and amperometric recordings of AMPH-induced efflux in isolated *Drosophila* brains were carried out as previously described (Campbell *et al.*, 2019).

### Statistics

GraphPad Prism 8.0 (GraphPad Software, San Diego, CA) software was used for data fitting and statistical analysis. Statistical methods for all comparisons are described in figure legends. Unless otherwise stated one-sample t-test was applied to examine relative differences i.e. for normalized data, while absolute measures were compared using either paired or unpaired t-test (depending on the experimental design) for normally distributed data or Mann-Whitney test for data that failed normality tests. One-way or two-way ANOVA with Holm-Sidak posttest were used for multiple comparisons.

### Study approval

Patient 1 and 2 were included from included from previously described patient samples (Fischbach & Lord, 2010; Hansen *et al.*, 2014). Informed consent was obtained was from patient 2 for all further investigations. The iPSYCH study sample is approved by the Danish Data Protection Agency. Informed consent is not required by law for register-based research in Denmark. Experimental procedures on animal adhered to the European guidelines for the care and use of laboratory animals, EU directive 2010/63/EU and were approved by the Danish Animal Experimentation Inspectorate (permission numbers 2017-15-0201-01160 and 2017-15-0201-01177). All efforts were made to minimize pain and discomfort as well as the number of animals used in each experiment.

## Acknowledgement

We are grateful to the patients and their relatives for participating in this study. We thank Anette Dencker Kaas, Annika H. Runegaard, Tina Skjørringe, and Pernille Emilie Petersen for excellent technical assistance. We thank the iPSYCH research leaders: Merete Nordentoft, Preben Bo Mortensen, Anders Børglum, Ole Mor, and David Hougaard along with Vivek Appadurai and Alfonso Demur for look-up in the iPSYCH database. We also own gratitude to Dominic Rizzi for the referral of patient 2 for clinical evaluation by LH. The work was supported by: Independent Research Fund Denmark – Medical Sciences (DFF-4183-00571, FH; DFF 4004-00097B, UG), Lundbeck Foundation R181-2014-3090 and R303-2018-3540, FH), Lundbeck Foundation (R223-2016-261, UG), Augustinus Fonden (11870, UG), National Institutes of Health (NIH) (DA035263, AG; HM), and (Z1A DA000610, AHN).

## Author contributions

FH and UG conceptualized the study. LEH identified, examined, and included patient 2 and coordinated his clinical procedures. Information on patient 1 was provided by AG. FH, KLJ, ST, NVA, AS, JA, LPP, AHR, HM, KE, TS, VKL and MR, conducted the in vivo, ex vivo, and in vitro experiments. AL and MNL provided and quantified DAT SPECT scans. HM and AG generated the untagged drosophila KI lines while OK and VKL generated the GFP-tagged drosophila lines, TRC and ATS designed and cloned AAV constructs. LBM and TS carried out sequencing of patient 2 and his parents. TW provided the iPSYCH information on DAT-K619N from exome data. AHN provided JHC 1-064 and MFZ 9-18, synthesized in the Medicinal Chemistry Section, NIDA-IRP. FH, KLJ, and UG prepared data and performed the formal data analyses. FH and UG provided funding, prepared figures and wrote the manuscript. All authors contributed to the editing and review of the manuscript.

## Competing interests

The authors have declared there are no conflicts of interest.

## Notes

### Competing Interest Statement

The authors have declared no competing interest.

