## Supplementary Material for "A dominant-negative variant in the dopamine transporter PDZ-binding motif is linked to parkinsonism and neuropsychiatric disease"

[1]Molecular Neuropharmacology and Genetics Laboratory, Department of Neuroscience, Faculty of Health and Medical Sciences, University of Copenhagen, Copenhagen, Denmark, [2]Department of Molecular Physiology & Biophysics, Vanderbilt University, Nashville, USA, [3]Department of Clinical Physiology & Nuclear Medicine, Bispebjerg Hospital, Copenhagen, Denmark. [4]National Institute on Drug Abuse-Intramural Research Program, National Institutes of Health, Baltimore, USA, [5]Department of Neurology, Bispebjerg Hospital, Copenhagen University Hospital, Denmark. [6]Institute of Biological Psychiatry, Mental Health Services Copenhagen; Department of Clinical Medicine, University of Copenhagen; and The Lundbeck Foundation Initiative for Integrative Psychiatric Research (iPSYCH), Denmark. [*]On behaf of iPSYCH researchers. [7]Center for Applied Human Genetics, Kennedy Center, Copenhagen University Hospital, Glostrup, Denmark [8]Department of Surgery, University of Alabama, Birmingham, USA, [9]Danish Dementia Research Centre, Clinic of Neurogenetics, Department of Neurology, Rigshospitalet, Copenhagen University Hospital and Department of Cellular and Molecular Medicine, Section of Neurogenetics, Faculty of Health and Medical Sciences, University of Copenhagen, Copenhagen, Denmark.

[†]Address Author correspondence to: Freja Herborg, Department of Neuroscience, Maersk Tower 7.5, University of Copenhagen, Blegdamsvej 3B, DK-2200 N, Copenhagen, Denmark. Phone +4553609699;. Or to: Ulrik Gether, Department of Neuroscience, Maersk Tower 7.5, University of Copenhagen, Blegdamsvej 3B, DK-2200 N, Copenhagen, Denmark. Phone +45 2875 7548;.

**Supplementary methods**

*Exome sequencing and variant quality control*

Samples were exome sequenced using the Illumina Nextera Rapid Capture kit at 20x average depth. The human reference genome version hg19 was used to align the sequenced reads, using the BWA short read aligner (Li & Durbin, 2009). Picard tools (https://broadinstitute.github.io/picard/) were applied to mark duplicates and the genome analysis tool-kit (GATK)'s HaplotypeCaller (McKenna *et al*, 2010) was used for variant calling. Variant quality control was performed using the Hapmap (The International HapMap *et al*, 2005) as truth and 1000 genomes variants as training datasets and carried out in accordance with the GATK's variant quality score recalibration modules. Variants that were outside the regions enriched as part of the exome capture kit design as well as monomorphic variants were excluded using Bedtools (Quinlan & Hall, 2010).

*Sample Quality Control*

The kinship coefficients within the sequenced samples were calculated using KING (Manichaikul *et al*, 2010), in case of samples showing higher than a third-degree relationship, the sample with higher genotyping call-rate was retained. The variants in the remaining samples were intersected with the variants present at a higher than 1% minor allele frequency (MAF) in the 1000 genomes phase3 call-set (The Genomes Project *et al*, 2015). The resulting variants common to the 1000 genomes and iPSYCH exome sequencing dataset were pruned using PLINK (Chang *et al*, 2015) with an LD window of 1mb and an $r^2$ threshold of 0.05. FlashPCA (Abraham & Inouye, 2014) was used to compute a principal component space using the resulting variants and treating the CEU (Central Europeans from Utah) population of the 1000 genomes as reference, an outlier detection algorithm, Aberrant (Bellenguez *et al*, 2012) was used to exclude iPSYCH samples where the inlier to outlier standard deviation ratio within the first two principal components exceeded 20:1. A total of 17339 samples passed sample QC.

**Supplementary Figures and Tables**

| | $K_M$ DA mean ± SEM (µM) | $V_{max}$ DA Mean ± SEM (% of WT) | $K_i$ CFT Mean (nM) [SEM interval] | $B_{max}$ CFT Mean ± SEM (% of WT) | AMPH IC50 (µM) [SEM interval] | MTP IC50 (nM) [SEM interval] | Cocaine IC50 (nM) [SEM interval] |
|---|---|---|---|---|---|---|---|
| WT | 0.84 ± 0.1 | 100 | 7.58 [6.3-9.3] | 100 | 1.03 [0.98-1.1] | 29.0 [26-33] | 122 [110-140] |
| K619N | 0.86 ± 0.1 | 71 ± 5* | 7.48 [6.4- 8.8] | 75.0 ± 5* | 1.11 [0.82-1.5] | 31.2 [30-32] | 118 [100-130] |

**Supplementary Table 1. DAT-K619N does not display altered interaction with ligands**. Kinetic parameters for $^3$H-dopamine uptake and $^3$H-CFT binding and IC50 values for classical substrates and inhibitors were evaluated in transiently transfected Cos-7 cells. Saturation [$^3$H]-dopamine uptake experiments were fitted to Michaelis Menten kinetics to derive $V_{max}$ and $K_M$ values, and nonlinear regression analysis of [$^3$H]-CFT/CFT competition binding curves was performed to calculate CFT Ki values and $B_{max}$. DAT-K619N showed significantly reduced maximal uptake and binding capacity ($p<0.05$, one sample t-test) without accompanying changes in $K_M$ or $K_i$ values. Ligand IC50 values were calculated from pIC50 values, determined by nonlinear regression analysis of competition [$^3$H]-dopamine uptake experiments. Mean CFT $K_i$ and ligand IC50 values with SE interval were calculated from the mean $pK_i$/pIC50 ± SEM and analysis for statistical differences to WT DAT was performed on the $pK_i$/pIC50 values. Data shown as mean ± SEM, and all experiments were performed in triplicates.

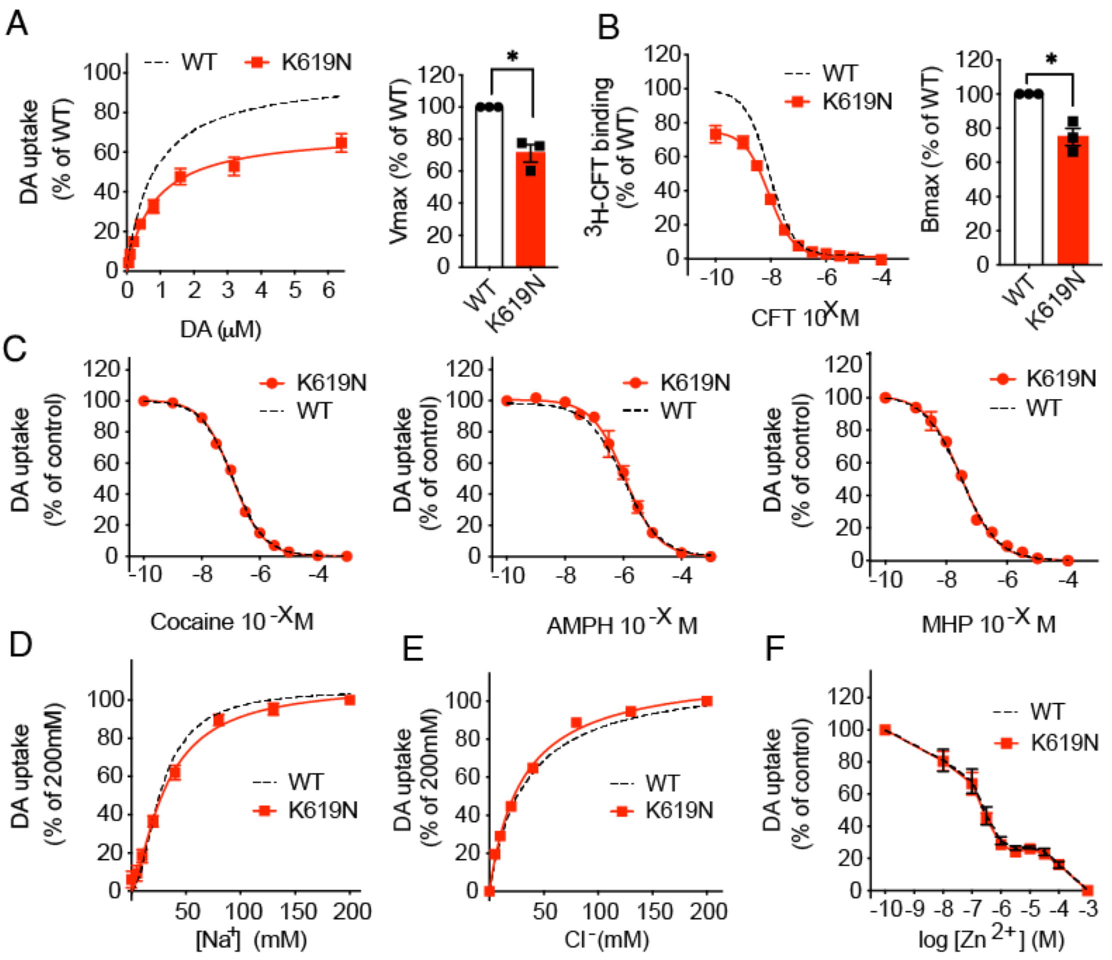


**Supplementary Figure 1. The DAT-K619N mutation does not disrupt interactions with ligands.** Ligand interactions, ion coordination and $Zn^{2+}$-dependent regulation of DAT-K619N in transiently transfected Cos-7 cells. [$^3$H]-DA uptake (A) and [$^3$H]-CFT/CFT competition binding experiments (B) on transiently transfected Cos-7 cells confirm the functional impairments of DAT-K619N observed in HEK329 cells ($p<0.05$, one-sample t-test, N=3). (C) Competition [$^3$H]- DA uptake curves for amphetamine, methylphenidate, and cocaine show no changes in apparent affinity for DAT-K619N. Data is normalized to control without inhibitor present ($p>0.05$, unpaired t-test analysis of IC50 values, N=3, see Supplementary Table 1). Kinetic parameters and IC50 values for A-C are listed in Supplementary Table 1. (D+E) Coordination of co-transported ions by DAT-K619N assessed by sodium (L) and chloride (M) dependency of [$^3$H]- DA uptake. Data is normalized to 200mM NaCl and equimolar cation and anion substitution were achieved with choline chloride and sodium gluconate respectively. No differences in ion coordination between DAT-K619N and DAT-WT were observed. (N) $Zn^{2+}$-dependent regulation of DA uptake by DAT-K619N is similar to DAT-WT, suggesting that the translocation cycle is not compromised by conformational changes. Curves show DA uptake normalized to control (absence $Zn^{2+}$). All curves are average curves of three independent experiments (mean ± SEM), each performed in triplicates. DAT-WT is presented as a dotted line as the experiments were conducted alongside a head-to-head comparison of previously reported disease-associated coding DAT variants published in (Herborg *et al*, 2018). *$p<0.05$, **$p<0.01$.

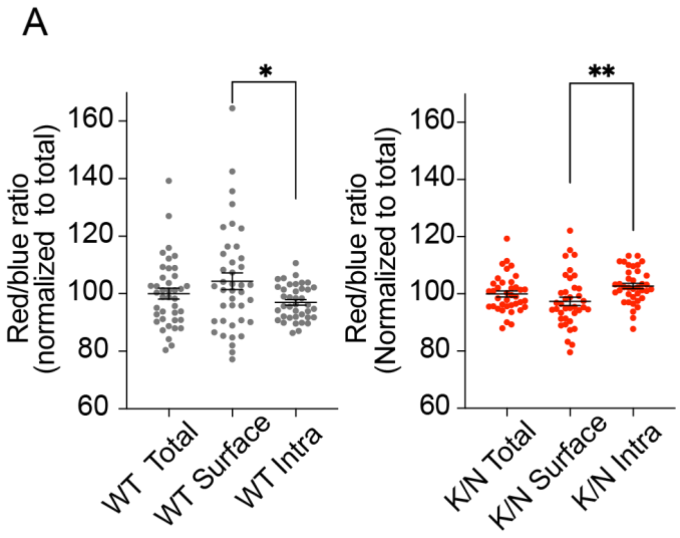


**Supplementary Figure 2. Fluorescent timer reveals changes in the DAT-K619N trafficking. (**A) Quantification of the red-to-blue ratios for the total protein pool, as well as the for the surface, and intracellular compartments separately show differences in the relative ages between SlowFT-DAT-WT and SlowFT-DAT-K619N. A green fluorescent cocaine analogue, MFZ 9-18, labelling DAT at the cell surface is used to separate the total visualized protein into a surface and an intracellular compartment. To compare the relative protein ages in the surface and intracellular compartments with that of the mean 'age' of the total protein pool, the red-to-blue ratios have been normalized to the mean age of the total protein for SlowFT-DAT-WT and SlowFT-DAT-K619N respectively. On average, SlowFT-DAT-WT at the cell surface is older than in the intracellular compartment (higher red to blue ratio). In contrast, the relative age for SlowFT-DAT-K619N is higher in the intracellular fraction than in the surface fraction ($p<0.05$, one-way ANOVA with Holm-Sidak correction for multiple comparison, N=36-39), indicating altered cellular processing of SlowFT-DAT-K619N. Data are shown as means ± SEM. Image analysis was done in ImageJ (see methods) *$p<0.05$, **$p<0.01$, ****$p<0.0001$.

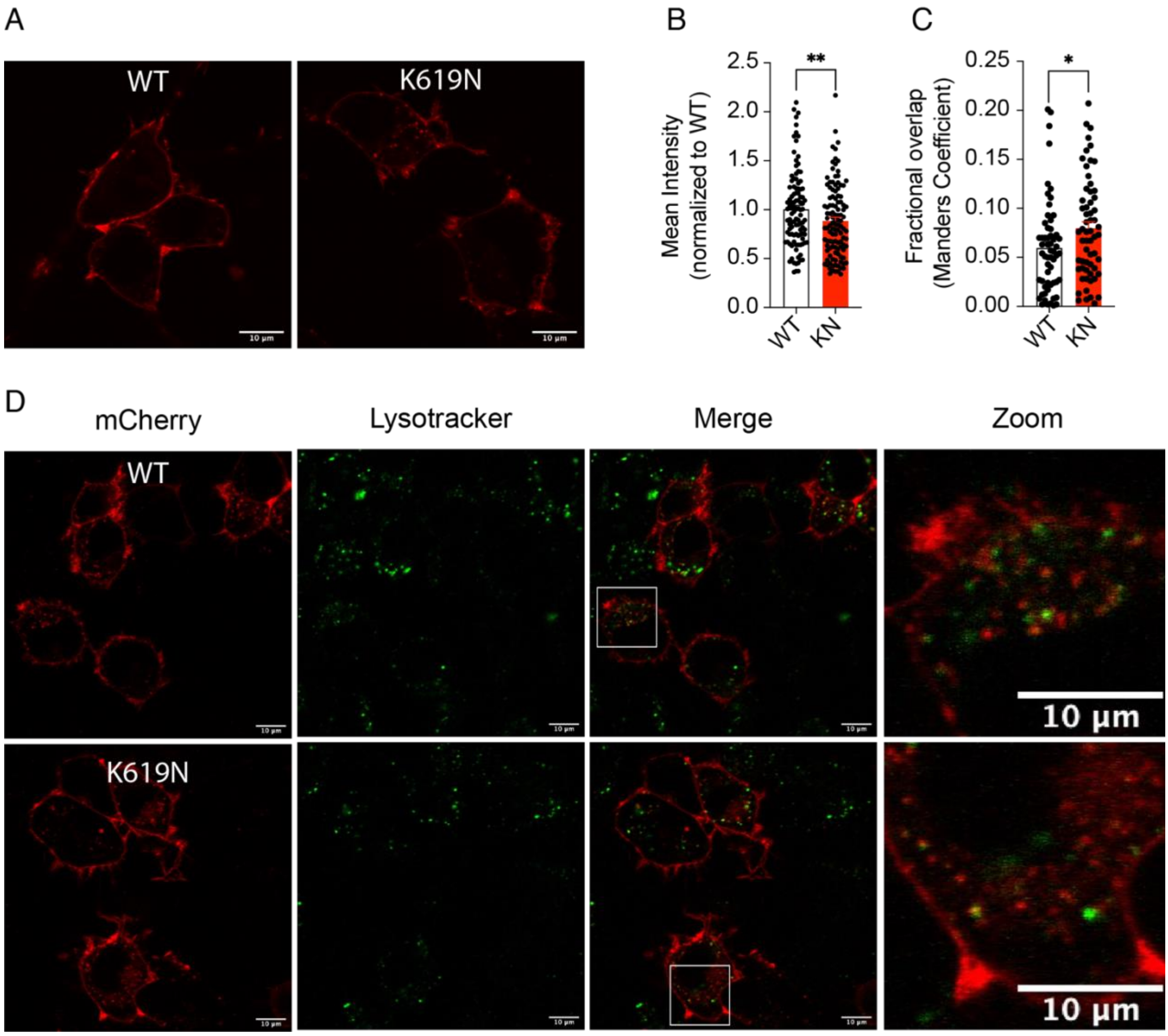


**Supplementary Figure 3. DAT- K619N demonstrates a larger fractional overlap with lysotracker.** (A) Live confocal microscopy of CAD cells transiently transfected with mCherry-DAT WT or mCherry-DAT-K619N. (B) Quantification of mean mCherry intensity. The mean mCherry intensity of all images was normalized to the WT average mean intensity for each imaging session. The relative mean intensity of mCherry-DAT-K619N is reduced compared to mCherry-DAT-WT (one-sample t-test, $p<0.01$, N=106-114 images). (C+D) Quantification and visualization of the fractional overlap between LysoTracker® Green and mCherry-DAT-WT or mCherry-DAT-K619N. Lysozymes were visualized by 15 min incubation with 50 nM LysoTracker® Green DND-26 probe before confocal live imaging. The relative amount of mCherry-DAT-WT and mCherry-DAT-K619N that colocalize with LysoTracker® positive structures was quantified using Manders coefficient (D) with the JaCoP Plug-in for ImageJ. mCherry-DAT-K619N showed a larger fractional overlap with lysosomes than mCherry-DAT-WT (fractional overlap=0.060±0.006 for DAT-WT vs 0.080±0.007 for DAT-K619N, $p<0.05$, Mann-Whitney test), consistent with enhanced lysosomal degradation. Images were acquired from at least three independent imaging sessions of at least three independent transfections. Representative images are shown. Scale bars are 10 µm.

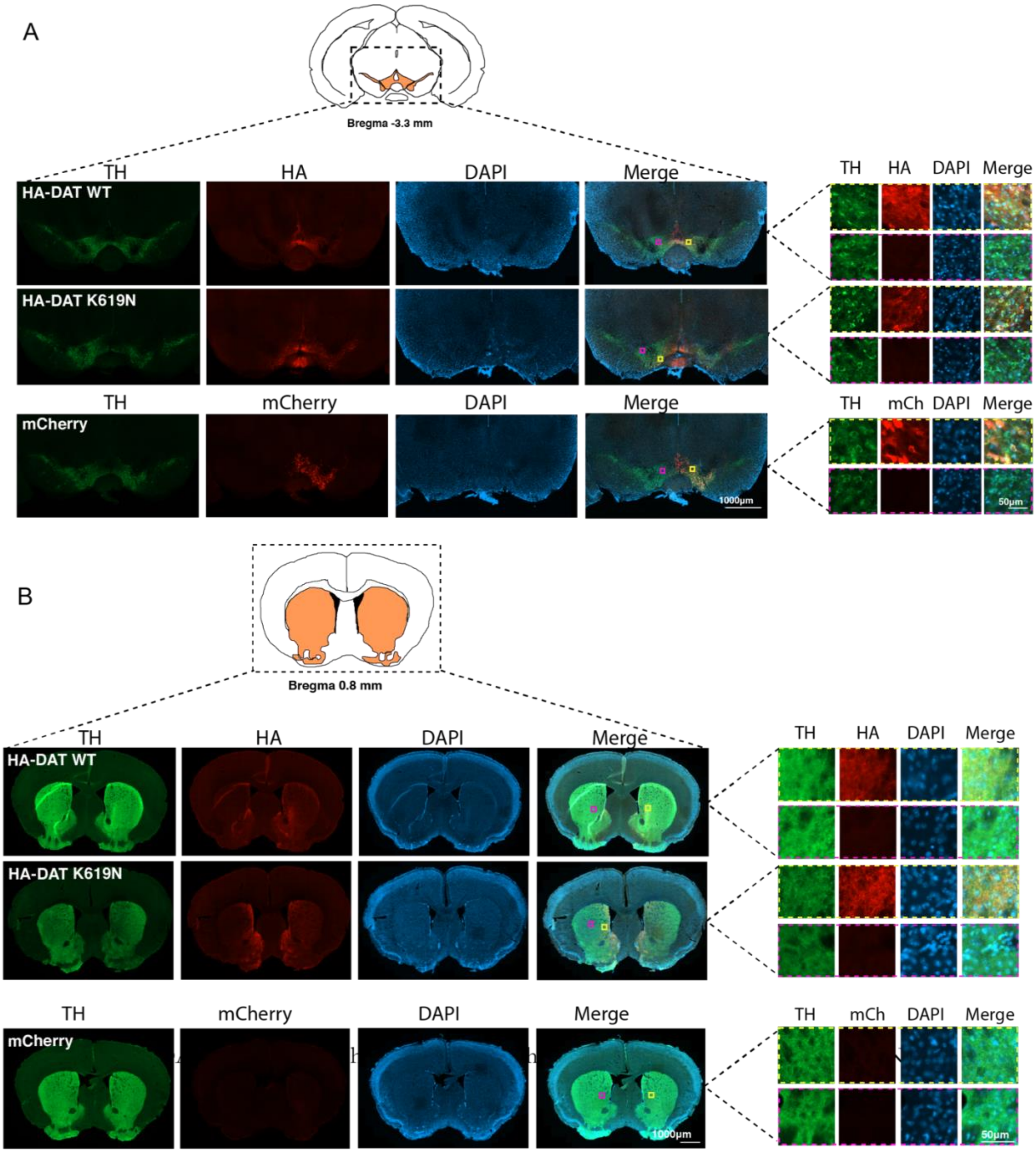


**Supplementary Figure 4. Viral expression of HA-DAT-WT and HA-DAT-K619N in midbrain dopaminergic neurons**. Immunohistochemical stainings of coronal slices from TH-cre mice injected in VTA with the following AAV constructs: DIO-hSYN-AAV8-HA-hDAT-WPRE, DIO-hSYN-AAV8-HA-hDAT-K619N-WPRE, or DIO-hSYN-AAV8-mCherry. Immunolabelling of midbrain (A) and striatal sections (B) against, TH, HA/mCherry and DAPI are shown. Mice injected with AAV encoding mCherry were not labelled for HA, but imaged for the exogenous mCherry expression, which is only found in the somatic region of midbrain dopaminergic neurons (A+B bottom panels). Magnifications inside and outside the viral infected area are shown in the right panels (N=2-3 mice of each genotype).

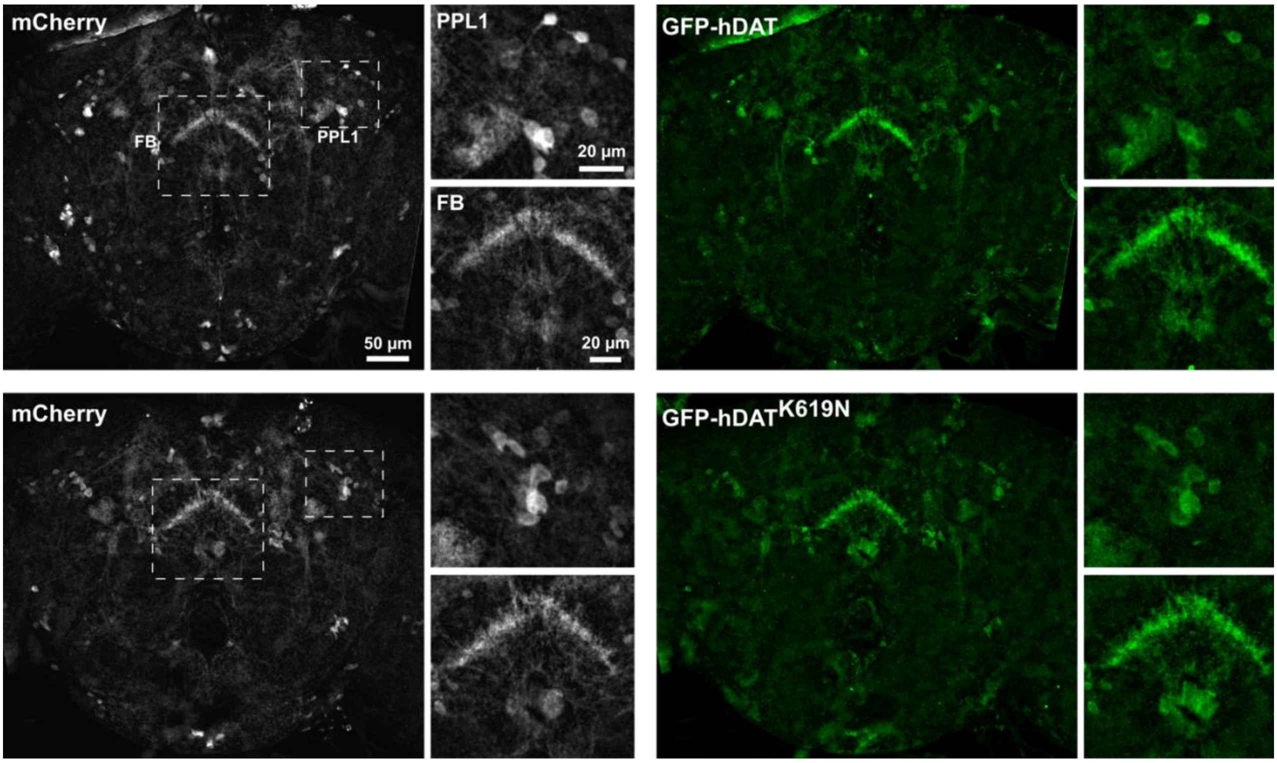


**Supplementary Figure 5. Visualization of EGFP-hDAT-WT and EGFP-hDAT-K619N in *Drosophila Melanogaster*.** (A) Confocal images of posterior brains from young adult *fmn* flies expressing either WT GFP-hDAT (*top*) or GFP-hDAT$^{K619N}$ (*bottom*) together with the mCD8-mCherry membrane marker in dopaminergic neurons, fixed and stained against GFP. Insets show dopaminergic axonal arbors in the fan shaped body (FB, *bottom*) and the dopaminergic paired posterior lateral 1 cell body cluster (PPL1, *top*) in high magnification. Images have been background subtracted. (B) Quantification of the GFP and mCherry signals from the FB arbor showed no difference in integrated intensity (P>0.05, unpaired t-test, N= 13)
